# Synergistic latency reversal by the IAP antagonist AZD5582 and BET inhibitor JQ1, combined with Nef ablation, facilitates immune-mediated elimination of latently HIV-1-infected T-cells

**DOI:** 10.1101/2025.08.06.668975

**Authors:** Dylan Postmus, Sui Ting Hui, Bengisu Akbil, Jenny Jansen, Dominic Wooding, Oya Cingöz, Christine Goffinet

## Abstract

The ‘shock-and-kill’ strategy can reactivate latent HIV-1 *in vivo* but has failed to meaningfully reduce the reactivatable reservoir, suggesting insufficient immunological visibility of infected cells. To investigate latency reversal agent (LRA)-specific shortcomings, we developed a novel HIV-1 latency model - Jurkat E6.1 subclonal T-cell lines, each harbouring a single, full-length, NL4.3-based provirus with a GFP_OPT_ reporter in the Env V5 loop - and quantified HIV-1 reactivation, Env surface presentation across LRAs and synergistically acting combinations. Bryostatin-1 induced GBP5-mediated restriction of surface Env and resistance to apoptosis. Bryostatin-1/JQ1 synergised for HIV-1 reactivation and boosted levels of cell-surface Env, but failed to reduce resistance to apoptosis. AZD5582/JQ1 boosted cell-surface Env levels, without co-induction of GBP5, T-cell activation or apoptotic resistance. HIV-1 Nef antagonised cytotoxic killing of HIV-1-reactivating T-cells - its knockout, combined with AZD5582/JQ1, achieved the highest immune-mediated elimination of latently infected T-cells, representing a promising HIV cure approach.

## INTRODUCTION

Recent rollbacks in international aid funding threaten the progress made globally in controlling and curbing the HIV pandemic, with the resulting disruptions in supply chains for antiretroviral therapy (ART) predicted to result in up to 10.75 million new infections in low- and middle-income countries between 2025 and 2030^1^. More than ever, these developments underscore the pressing need to establish an accessible, scalable and affordable cure strategy for people living with HIV (PLWH).

To this end, the ‘shock-and-kill’ cure strategy proposes the transcriptional reactivation of the latent provirus, with the aim of rendering infected cells immunologically visible, a key requirement for cytotoxic killing by patrolling immune cells. While conceptually promising, several in-human clinical trials attempting this intervention have fallen short of the desired outcome. Notably, the ROADMAP^2^ and eCLEAR^3^ trials, both of which administered the histone deacetylase inhibitor (HDACi) romidepsin, in combination with the broadly neutralising, CD4 binding site-targeting, anti-HIV-1 Envelope (Env) antibody 3BNC117, showed varying efficacies in delaying viral rebound post-treatment interruption. The ROADMAP trial demonstrated significant but modest increases in cell-associated unspliced HIV RNA (1.14 median fold increase) and detectable viral RNA copies in serum for the romidepsin/3BNC117 combination-treated group. The eCLEAR trial, which administered this intervention in newly diagnosed PLWH shortly after ART initiation, failed to report a statistically significantly enhanced reduction of the size of the viral reservoir as compared to the decay detected after treatment with ART alone. Additional knowledge on ‘shock-and-kill’ approaches has been generated in animal models of HIV-1 infection, such as HIV-1-infected humanised mice and SIV-infected non-human primates (NHPs), using LRAs of different classes such as inhibitors of inhibitor of apoptosis proteins (iIAPs)^4^ and inhibitors of bromodomain and extraterminal motif proteins (iBETs)^5^. Importantly, a landmark study in NHPs identified the combination of the iIAP AZD5582 with SIV-specific anti-Env antibodies was able to reduce, significantly, the number of intact proviruses in lymph nodes, but not the peripheral blood or spleens of ART-treated rhesus macaques^6^. While this approach showed promise, it did not achieve functional cure. Collectively, the continuous inability of ‘shock-and-kill’ attempts *in vivo* to achieve cure likely reflects our lack of detailed understanding of the pro- and antiviral impacts individual LRAs may exert in the context of latency reversal, and indicate a requirement for a more comprehensive understanding on the cellular effects of these treatments. In cellular studies, a myriad of different LRAs and LRA combinations have shown, with varying efficiencies and often in simplistic experimental models, ability to reactivate viral transcription and, to some degree, expression of a reporter and/or of viral antigens^7^. However, their impact on subsequent processes crucial for immune clearance, including peptide and antigen presentation, and susceptibility to cytotoxic killing, has been less attentively analysed. Here, we compared two different synergistically-acting LRA combinations - the iBET JQ1, paired with either bryostatin-1 (Bryo), a protein kinase C agonist (PKCa) and activator of canonical NF-κB signalling, or AZD5582 (AZD), an iIAP and activator of non-canonical NF-κB signalling - as potential ‘shock-and-kill’ interventions. While both strategies shared the ability to potently reverse HIV-1 latency, as previously described^8,9^, they vastly differed regarding induction of cell-surface accessibility of the viral Env protein and modulation of the cellś susceptibility to cytotoxic killing. Finally, LRA treatment-mediated co-induction of Nef expression antagonised increased cellular resistance to killing by almost all tested latency reversal strategies, advocating for the integration of a Nef-targeting approach into future ‘shock-and-kill strategy’ development.

## RESULTS

### Bryostatin-1, despite its latency reversal activity, imposes a restriction to HIV-1 Env cell-surface presentation

In addition to presentation of peptides by MHC-I, accessibility of the HIV-1 Env glycoprotein on the cell surface for binding to anti-Env antibodies is crucial for immunological visibility of HIV-1-reactivating cells. We assessed how different classes and combinations of LRAs modulate HIV-1 Env expression and subcellular localisation, as well as cellular vulnerability to cytotoxic killing. We selected bryostatin-1 and JQ1 as prototypical PKC agonist and BET inhibitor, respectively, because they have been described to act synergistically^8^. The HDAC inhibitor panobinostat (Pano) was included as an additional class of clinically relevant, widely used LRA^10,11^. In this study, we used Jurkat E6.1 T-cells, latently infected with a full-length, replication-competent HIV-1 NL4.3 reporter virus bearing an optimised GFP sequence (GFP_OPT_) fused into the V5 loop domain of gp120 (Figure S1A)^12^. Three individual, clonal cell lines of Jurkat T-cells, each carrying a single latent provirus per cell with known sequence and integration site (Jurkat Latent EnvGFP [JLEG]: JLEG.1, JLEG.60 and JLEG.74, Table S1), expressed low-to-undetectable p24 capsid protein (p24CA) and EnvGFP in the absence of stimulation (Figure 1A). Upon treatment with individual LRAs, including bryostatin-1, JQ1, panobinostat, and, most pronouncedly, a combination of bryostatin-1 and JQ1, JLEG.1 T-cells expressed detectable p24CA, EnvGFP, and scored positive for cell-surface EnvGFP as assessed by anti-GFP immunostaining of intact cells (Figure 1A, S2A). Matrix titrations of increasing concentrations of bryostatin-1 and JQ1, respectively, confirmed the previously described synergy of these two molecules’ ability to reverse HIV-1 latency^8^ in a Bliss-Independence test (Figure S1B).

**Figure 1.**
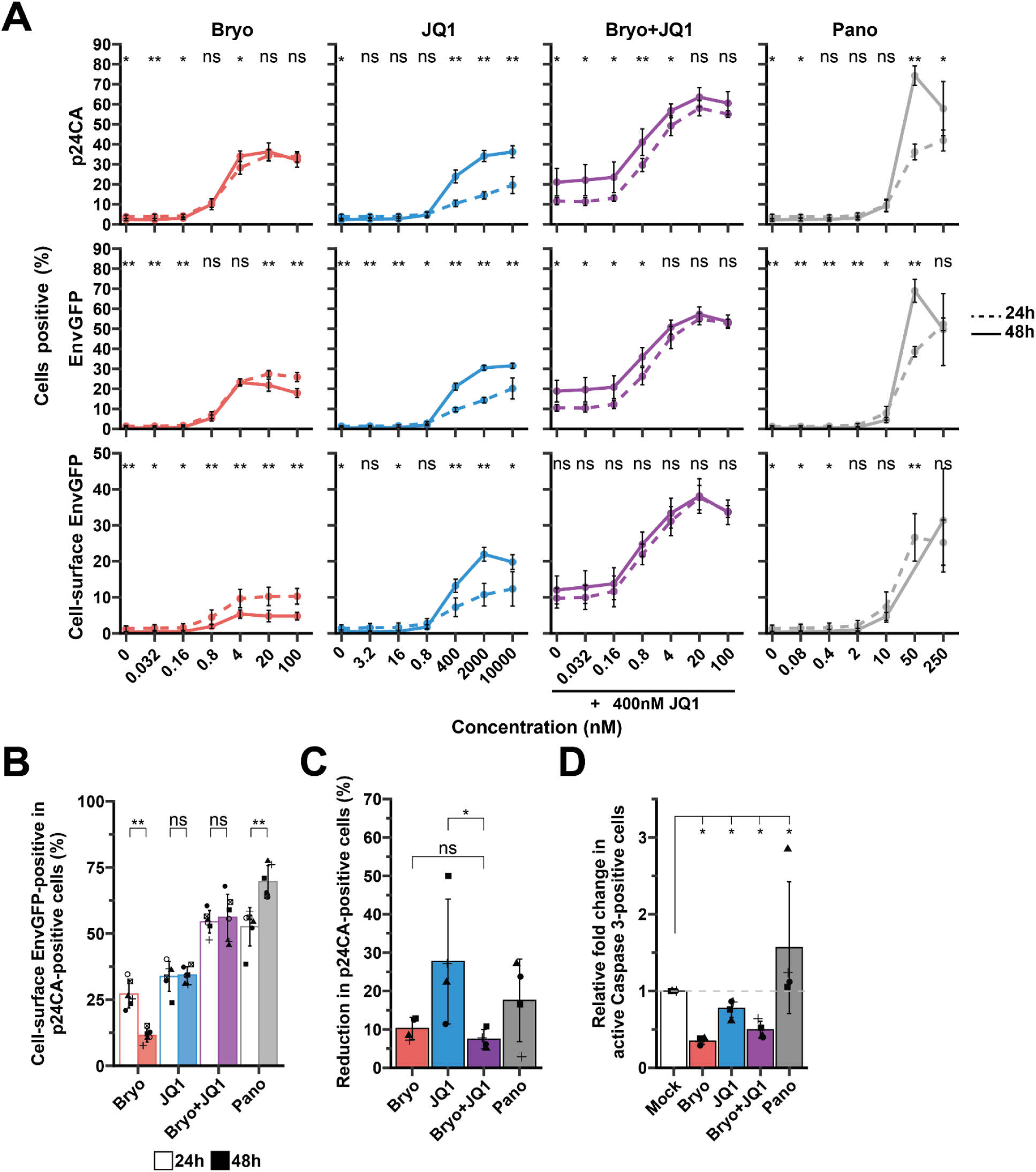
Bryostatin-1, despite its latency reversal activity, imposes a restriction to HIV-1 Env cell-surface presentation A) Titration of bryostatin-1, JQ1, bryostatin-1 in the presence of 400nM JQ1, and panobinostat on JLEG.1 cells, measuring intracellular p24CA, total EnvGFP and cell-surface levels of EnvGFP at 24h and 48h post-reactivation by flow cytometry; n=6, Wilcoxon rank-signed test. B) Quantification of cell-surface EnvGFP-positive cells within the p24CA-positive population in JLEG.1 cells treated for 24h and 48h with bryostatin-1 (4nM), JQ1 (400nM), a combination of bryostatin-1 and JQ1, or panobinostat (50nM), measured by flow cytometry; n=6xlink:href=" Wilcoxon rank-signed test. C) Reduction in p24CA-positive cells for JLEG.1 T-cells treated with the indicated LRAs for 24h, followed by co-culture with PBMCs from HIV-negative donors in the presence of sera from PLWH for 4h to induce antibody-dependent cytotoxic killing; n=4 PBMC donors, t-test on log-normalised data. D) Relative proportion of JLEG.1 T-cells expressing cleaved, active Caspase 3 upon indicated treatments for 24h, followed by exposure to etoposide (16h) to induce intrinsic apoptosis, measured by flow cytometry. Expression was normalised to the mock condition; n=4, Wilcoxon rank-signed test. *** p < 0.001; ** p < 0.01; * p< 0.05. Indicated is the arithmetic mean ± SD.

JQ1 concentrations of 400 nM and above resulted in a significant increase from 24 to 48 hours for all three parameters measured (Figure 1A). In contrast, while bryostatin-1 treatment induced appreciable levels of p24CA and total EnvGFP, the levels of cell-surface EnvGFP were relatively low and even decreased from 24h to 48 hours post-treatment. Co-treatment with JQ1 appeared to reverse or prevent this downregulation (Figure 1A, S2A-C). Quantification of the percentage of cell-surface EnvGFP-positive cells among p24CA-positive cells (Figure S2B) confirmed that among the tested LRA strategies, only bryostatin-1 mono-treatment interfered with cell-surface EnvGFP presentation over time (Figure 1B, Figure S2C). Despite displaying higher cell-surface EnvGFP levels compared to the respective mono-treatments, T-cells co-treated with bryostatin-1 and JQ1 showed the highest resistance to killing and poor overall clearance of infected cells in an antibody dependent cellular cytotoxicity (ADCC) assay (Figure 1C). This suggests that the combined treatment imposed a barrier to cytotoxic elimination of infected cells that was not overcome by increased availability of the Env antigen. Indeed, bryostatin-1 and JQ1 co-treatment induced a marked resistance to etoposide-mediated induction of intrinsic apoptosis (Figure 1D), which we hypothesised underlies the inefficiency of ADCC-mediated killing.

### Bryostatin-1 treatment induces expression of T-cell activation markers and GBPs

In order to gain deeper insight into alterations of the cellular state that might underlie the observed phenotypes, we performed total RNA sequencing (RNAseq) on JLEG.1 T-cells and an uninfected cellular clone from the same original Jurkat culture. We noted sizable and treatment-specific perturbations to the transcriptome, corresponding to high numbers of significantly up- and downregulated genes (Figure 2A-B) and a relatively large overlap in differentially regulated genes for the LRA treatments relative to the mock condition (Figure 2C). Importantly, the presence of a provirus also considerably shaped transcriptomic responses to individual treatment strategies (Figure 2D). Pathway enrichment analysis^13^ revealed that bryostatin-1 increased activity for pathways related to T-cell activation and effector phenotypes, including ‘Interferon gamma signalling’ (R-HSA-877300) and ‘Costimulation by the CD28 family’ (R-HSA-389356), and this was markedly diminished by addition of JQ1 (Figure 2E). Interestingly, analysis of the interaction effect between the presence of a provirus and LRA addition indicated cooperation between bryostatin-1 administration and viral reactivation to boost cholesterol biosynthesis pathways (Figure 2E), possibly relating to previous observations on the influence of HIV proteins on lipid metabolism^14–16^.

**Figure 2.**
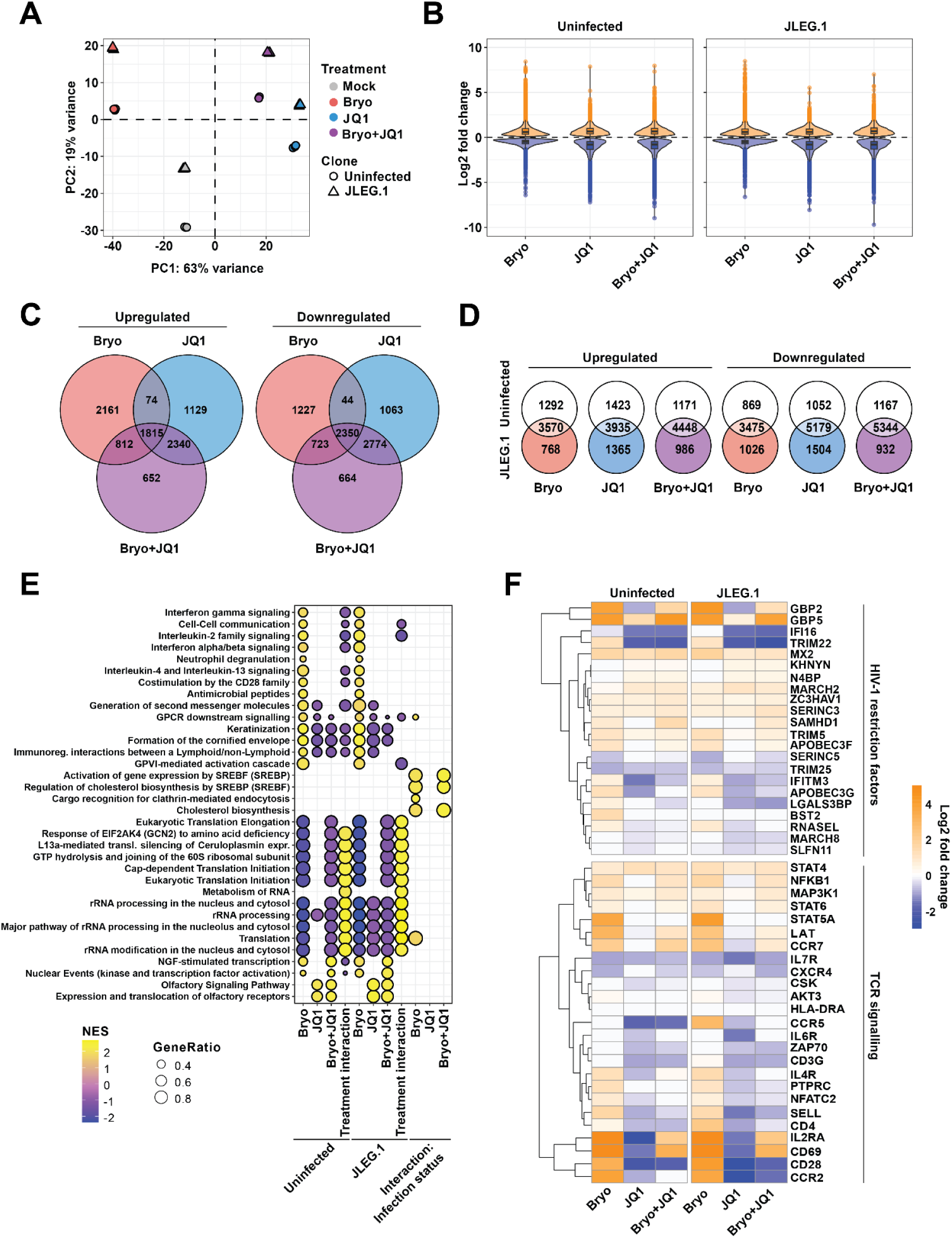
**Bryostatin-1 treatment induces expression of T-cell activation markers and GBPs.** A) Principal component analysis (PCA) plot of the transcriptomes of the JLEG.1 clone, as well as of an uninfected clone of same origin, after indicated treatments for 24h; n=3 for each clone and treatment combination. B) Violin plots indicating the statistically significantly up- (orange) and downregulated (blue) genes in the indicated treatments, relative to the mock condition (p_adjusted_ < 0.05). C) Venn diagrams illustrating the overlap in statistically significantly differentially up- and downregulated genes relative to the mock condition in JLEG.1 cells between treatments (p_adjusted_ < 0.05). D) Venn diagrams illustrating the overlap in statistically significantly differentially up- and downregulated genes relative to the mock condition for the indicated treatments between the uninfected clone (top) and JLEG.1 cells (bottom) (p_adjusted_ < 0.05). E) Gene set enrichment analysis of differentially upregulated genes for each treatment relative to the mock condition by respective cell line using the Gene Ontology: Biological Pathway database. Indicated are the top 20 enriched pathways by treatment and cell line. Indicated is also the interaction effect, representing the effect on the enrichment of certain pathways that is modulated by the presence of a provirus. F) Fold change in gene expression for each LRA treatment relative to the mock condition, determined for the uninfected and JLEG.1 clone respectively, for selected genes corresponding to known HIV-1 restriction factors and TCR signalling pathways.

When assessing the response by curated gene sets, treatment with bryostatin-1, either alone or in combination with JQ1, but not JQ1 alone, markedly upregulated T-cell activation markers *CD69* and *IL2RA*/*CD25^17,18^* (Figure 2F). In addition, bryostatin-1, both in the absence and presence of JQ1, significantly induced *GBP2* and *GBP5* RNA levels, restriction factors previously described to interfere with HIV-1 Env biosynthesis in a saturable manner^19,20^. In fact, *GBP5* was identified as one of the most highly upregulated genes, both when considering global transcriptomic changes across treatments (Figure S3A), and expression patterns within curated gene sets of HIV-1 restriction factors (Figure 2F) and interferon signalling (Figure S3B). We hypothesised that bryostatin-1 treatment, by driving T-cell activation, renders cells more resistant to apoptosis, while simultaneously limiting the levels of Env presentation via the co-induction of restriction factors targeting this viral protein. This would, in turn, make reactivated cells at once less immunologically visible and more resistant to apoptosis-inducing cytotoxic effector molecules.

### Bryostatin-1 treatment and reactivation of proviral transcription, but not AZD5582 treatment, promote T-cell activation

As bryostatin-1 appeared to induce a phenotype counterproductive to immune-mediated killing of infected cells, we decided to investigate the iIAP AZD5582 as a member of a different class of molecules shown to synergise with BET inhibitors^9^. In contrast to AZD5582 mono-treatment, treatment of all three JLEG T-cell clones with AZD5582 and JQ1 potently induced HIV-1 latency reversal and high levels of cell-surface Env to an extent that was comparable to the bryostatin-1 and JQ1 combination (Figure 3A, S4A). The effect was also confirmed to be synergistic under the Bliss-Independence model (Figure S4B).

**Figure 3.**
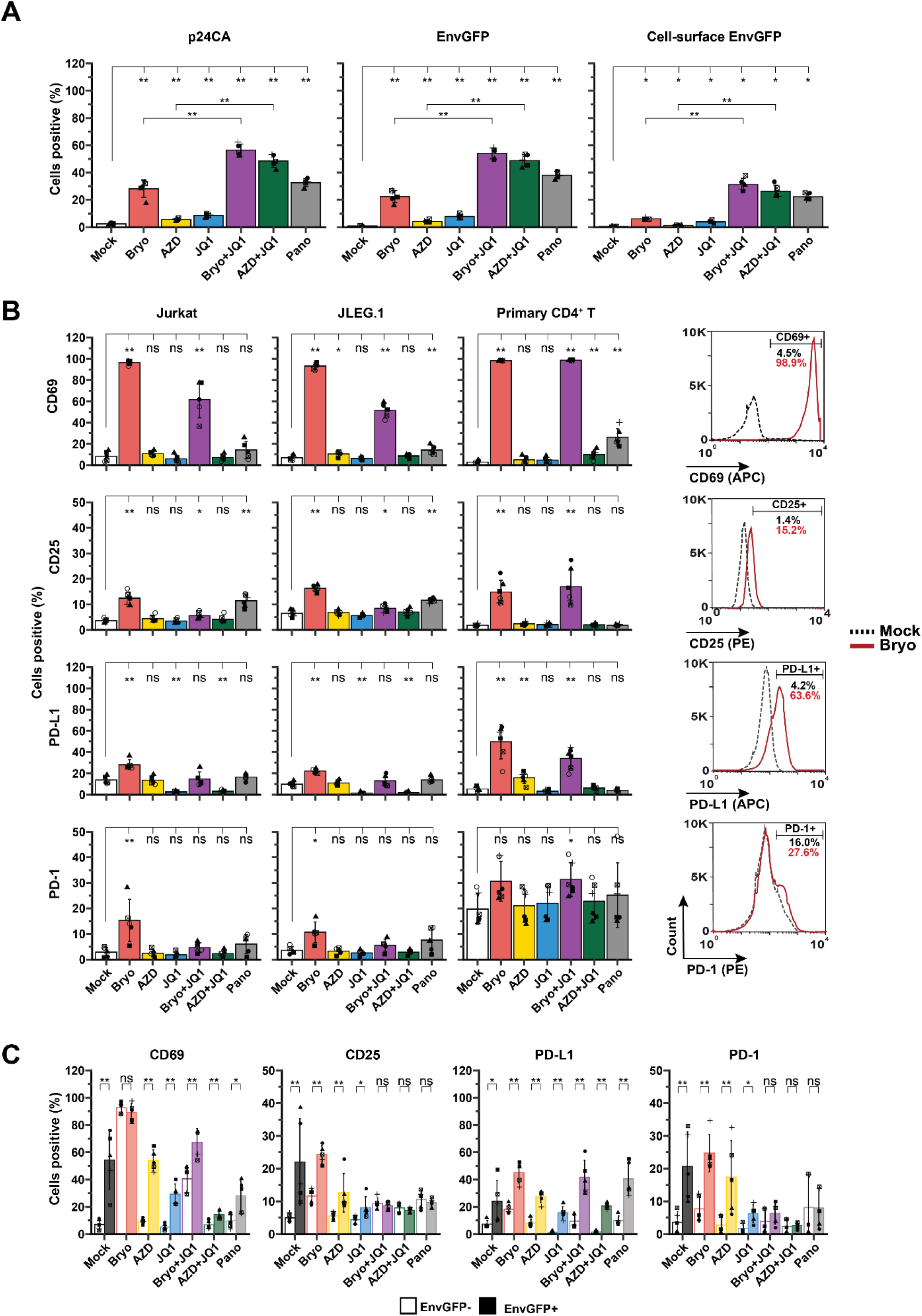
**Bryostatin-1 treatment and reactivation of proviral transcription, but not AZD5582 treatment, promote T-cell activation** A) Percentage of cells expressing HIV-1 p24CA (left panel), total EnvGFP (middle panel) and cell-surface EnvGFP (right panel) in JLEG.1 T-cells treated as indicated for 24h, quantified by flow cytometry; n=5. B) Percentage of cells positive for cell-surface expression of T-cell activation markers CD25, CD69, PD-L1 and PD-1 in JLEG.1; uninfected Jurkat T-cells; and unstimulated, primary CD4^+^ T-cells from HIV-1-negative donors treated as indicated for 24h, quantified by flow cytometry; n=6. Representative flow cytometry histograms for mock-treated and bryostatin-1-treated primary CD4^+^ T-cells are provided. C) Expression of T-cell activation markers on JLEG.1 T-cells, treated as above, assessed for EnvGFP-positive and EnvGFP-negative T-cells, respectively, in the same sample and quantified by flow cytometry; n=6. Wilcoxon rank-signed test. *** p < 0.001; ** p < 0.01; * p< 0.05. Indicated is the arithmetic mean ± SD.

As RNAseq analysis suggested that bryostatin-1 induces T-cell activation (Figure 2E-F), we sought to validate this on the protein expression level, while simultaneously investigating how coincident proviral reactivation shapes this response. Expectedly, treatment with bryostatin-1 and the bryostatin-1 and JQ1 combination upregulated expression of the early and early-intermediate activation markers CD69 and CD25 (IL2RA) in both the uninfected, parental Jurkat T-cells and JLEG.1 T-cells (Figure 3B). Bryostatin-1 mono-treatment also induced the expression of the late activation marker PD-L1 and the exhaustion marker PD-1, though this was dampened in these systems by addition of JQ1. Induction of these activation markers was almost or completely absent in treatment with AZD5582 or the AZD5582 and JQ1 combination. Importantly, the activation phenotype induced by bryostatin-1 and the bryostatin-1 and JQ1 combination but not by the AZD5582 and JQ1 combination was recapitulated in primary CD4^+^ T-cells from HIV-1-negative donors in that CD25, CD69, PD-L1 and PD-1 were markedly upregulated. As PD-L1 protects from CTL-mediated killing^21^, reactivation with bryostatin-1 could be profoundly counterproductive to a ‘shock-and-kill’ cure strategy from the perspective of cytotoxic T-cell efficacy. Panobinostat, in contrast, moderately induced CD69 in JLEG.1 and primary CD4^+^ T-cells and CD25 only in the Jurkat T-cell-based systems, without affecting PD-1 or PD-L1 expression. In line with a previous report in uninfected Jurkat T-cells^22^, bryostatin-1 administration reduced cell-surface levels of the HIV-1 receptor CD4, without altering expression of CCR5 in all T-cell models tested (Figure S5A), an observation analogous to what has previously been reported in macrophages^23^ and is in line with an activated T-cell phenotype^24^. All tested LRAs, with the exception of AZD5582 in primary CD4^+^ T cells, reduced the expression of CXCR4, and this was most striking in the bryostatin-1- containing treatments (Figure S5A). HIV-1-reactivating cells, expectedly^25–29^, had lower levels of cell-surface CD4 than non-reactivating cells, except in the case of bryostatin-1 containing treatments where the LRA had already induced significant downregulation of CD4. However, the described virus-mediated downregulation of CCR5^30,31^ was not observed, owing possibly to low overall levels of CCR5 expression in this system (Figure S5B). Interestingly, HIV-1 reactivating cells also downregulated cell-surface CXCR4 in accordance with previous reports^32,33^ for all treatments except for the bryostatin-1 mono-treatment, where reactivating cells displayed higher CXCR4 expression than non-reactivating cells, though the reason for this is unclear.

Discriminating between EnvGFP-negative (non-HIV-1-reactivating) versus EnvGFP-positive (HIV-1-reactivating) JLEG.1 T-cells in the same sample revealed a tendency for reactivating cells to display higher levels of activation markers (Figure 3C). This was true of PD-L1 in all treatments, including the mock condition, as well as CD69 (excepting the bryostatin-1 treatment, in which it was already maximally induced). CD25 and PD-1, additionally, were more highly displayed on reactivating cells in the mock, bryostatin-1, AZD5582 and JQ1 conditions (Figure 3C). This implies that not only treatment with bryostatin-1 and the bryostatin-1 and JQ1 combination, but also proviral reactivation itself, can drive T-cell activation programs.

Taken together, these data imply that the presence of the reactivating provirus *per se* can drive a T-cell activation phenotype to an extent shaped partially by the LRA administered. Additionally, they establish that the combination of AZD5582 and JQ1, as opposed to bryostatin-1 and JQ1, at the same time enhances immunological visibility of reactivating cells by supporting expression of high levels of cell-surface Env and avoids induction of an anti-apoptotic T-cell activation response that would otherwise be deleterious to immune clearance.

### GBP5 expression, induced by both bryostatin-1 treatment and reactivation of proviral transcription, is accompanied by reduced cell-surface levels of EnvGFP

RNA-seq analysis provided first evidence of the bryostatin-1 treatment inducing high levels of *GBP5* mRNA (Figure 2F). Given its well-documented ability to interfere with Env processing and trafficking^19,20,34^, we surmised that GBP5 induction may underpin the decrease in cell-surface Env levels at 48 hours relative to 24 hours post-treatment with bryostatin-1 (Figure 1A-B, Figure S2). Importantly, only bryostatin-1 and a reference PMA/ionomycin (PMA/Io) treatment, and not AZD5582, JQ1, or the combination of these showed a reduction of Env presentation over time (Figure S6A). Intracellular immunostaining for GBP5 (Figure 4A-B) confirmed the predicted upregulation of GBP5 upon treatment with bryostatin-1, even at sub-nanomolar concentrations. GBP5 induction also occurred with PMA/ionomycin and, to a lesser extent, during bryostatin-1 and JQ1 co-treatment, but not for any combination of AZD5582 and JQ1. These effects were replicated in primary CD4^+^ T cells, except in the case of panobinostat which only induced GBP5 in JLEG.1 T-cells (Figure 4B-D). Given that PMA/ionomycin is a known inducer of T-cell activation, the data suggest that GBP5 might be upregulated as a direct consequence of the activation state. Supporting this notion, other T-cell activation-inducing treatments commonly used *ex vivo* similarly upregulated GBP5 in primary CD4^+^ T-cells (Figure S6B).

**Figure 4.**
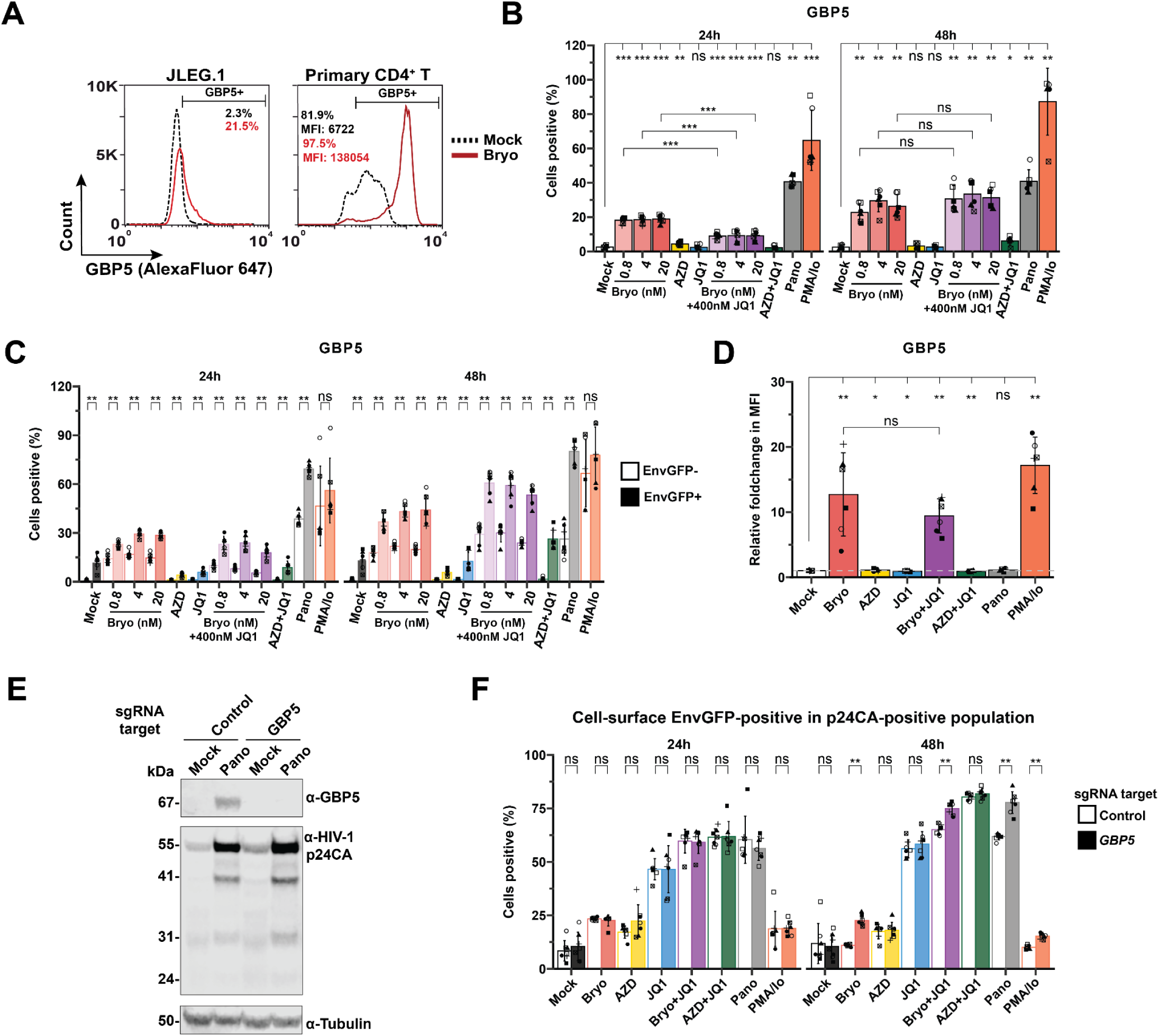
**GBP5 expression, induced by bryostatin-1 treatment and the reactivating provirus, is accompanied by reduced cell-surface levels of EnvGFP** A) Representative flow cytometry histograms illustrating the increase in GBP5 expression, indicated by percentage of GBP5-positive cells or the mean fluorescence intensity (MFI) of the GBP5 signal for JLEG.1 (left) and primary CD4^+^ T-cells (right) after mock treatment and bryostatin-1 treatment for 24h. B) Percentage of GBP5-positive JLEG.1 T-cells after mock treatment or treatment with the indicated LRAs, including varying, indicated concentrations of bryostatin-1, for 24h and 48h, respectively, quantified by flow cytometry; n=5-7. C) Percentage GBP5-positive JLEG.1 T-cells, within EnvGFP-negative and EnvGFP-positive cells of the same sample, treated as in A) for 24h and 48h; n=5-6. D) Fold change of GBP5 MFI, relative to the mock-treated condition, for primary CD4^+^ T-cells treated with the indicated LRAs. E) Representative immunoblot showing the CRISPR/Cas9-mediated knockout of the *GBP5* gene in JLEG.1 T-cells, either mock-treated or treated with panobinostat as a known inducer of GBP5 for 24h. The viral Gag proteins (detected by anti-p24CA antibody) are included as a control for reactivation efficacy. Tubulin is included as a loading control.. F) Expression of cell-surface EnvGFP in the HIV-1 p24 capsid-positive population for *GBP5* knockout or control JLEG.1 T-cells after mock treatment or treatment with the indicated LRAs for 24h and 48h. Wilcoxon rank-signed test. *** p < 0.001; ** p < 0.01; * p< 0.05. Indicated is the arithmetic mean ± SD.

Interestingly, HIV-1-reactivating JLEG.1 T-cells produced significantly higher levels of GBP5 than non-HIV-1-reactivating T-cells from the same culture for most LRA strategies tested, and importantly also under mock conditions (Figure 4C), indicating that viral reactivation may also, directly or indirectly, drive GBP5 expression. GBP2 was induced by bryostatin-1 treatment, both in JLEG.1 and primary CD4^+^ T-cells, though the relative increase was more modest than GBP5 (Figure S6C-D). Additionally, we noted that the boosted cell-surface Env levels in the bryostatin-1 and JQ1 combination condition (Figure S6A) persisted in spite of coincident GBP5 upregulation (Figure 2F, 4B-D). We concluded that this likely relates to the excess of EnvGFP overcoming any meaningful restriction by GBP5^34^.

Importantly, *GBP5* knockout in the JLEG T-cell lines (Figure 4E-F, Figure S6E-F) resulted in significantly higher cell-surface Env levels for all LRA regimens that induce GBP5 at 48 hours post-treatment, mechanistically implicating GBP5 upregulation in the observed cell-surface Env downregulation. This tendency for bryostatin-1 to limit Env antigen display via coincident restriction factor upregulation is problematic for antibody-mediated detection of infected cells and brings into question the usefulness of this LRA for facilitating immune-mediated clearance of the viral reservoir.

### Bryostatin-1 and the reactivating provirus both antagonise induction of apoptosis

While ADCC is initiated by effector cell recognition of antibody bound to a cell-surface antigen (Env, in this case) expressed on target cells, the subsequent cytotoxic killing is mediated largely by secretion of effector molecules that induce intrinsic and extrinsic apoptosis, such as Granzyme B and Fas (CD95) ligand (FasL), respectively^35^. The relatively low ADCC efficiency (Figure 1D) despite high cell-surface Env expression (Figure 1B-C), as well as the apparent reduced sensitivity to intrinsic apoptosis observed in latently infected T-cells treated with the bryostatin-1 and JQ1 combination (Figure 1E), suggests that impaired apoptosis signalling is a major contributor to the observed resistance to killing. In uninfected Jurkat T-cells, only bryostatin-1-comprising treatments limited intrinsic apoptosis, compared to the mock treatment, while all LRA regimens tested preserved the susceptibility of parental Jurkat cells to extrinsic apoptosis (Figure 5A). In stark contrast, HIV-1-infected JLEG.1 T-cells displayed resistance to cell death by both intrinsic and extrinsic apoptosis upon reactivation with all treatments except AZD5582 or panobinostat mono-treatment, providing a first hint for virus-mediated antagonism of apoptosis induction (Figure 5A). Indeed, HIV-1-reactivating T-cells were markedly more resistant to both routes of apoptosis than non-reactivating T-cells in the same sample (Figure 5B), indicating that one or several viral gene products can counteract apoptosis in this system. Importantly, panobinostat treatment alone, also when compared to the other LRA regimens, induced a high level of apoptosis, a phenomenon described previously^36^ and noticeably absent in primary CD4^+^ T-cells (Figure S7A). This level of pre-existing apoptosis prior to addition of any pro-apoptotic stimuli may make the results concerning apoptosis susceptibility more difficult to interpret.

**Figure 5.**
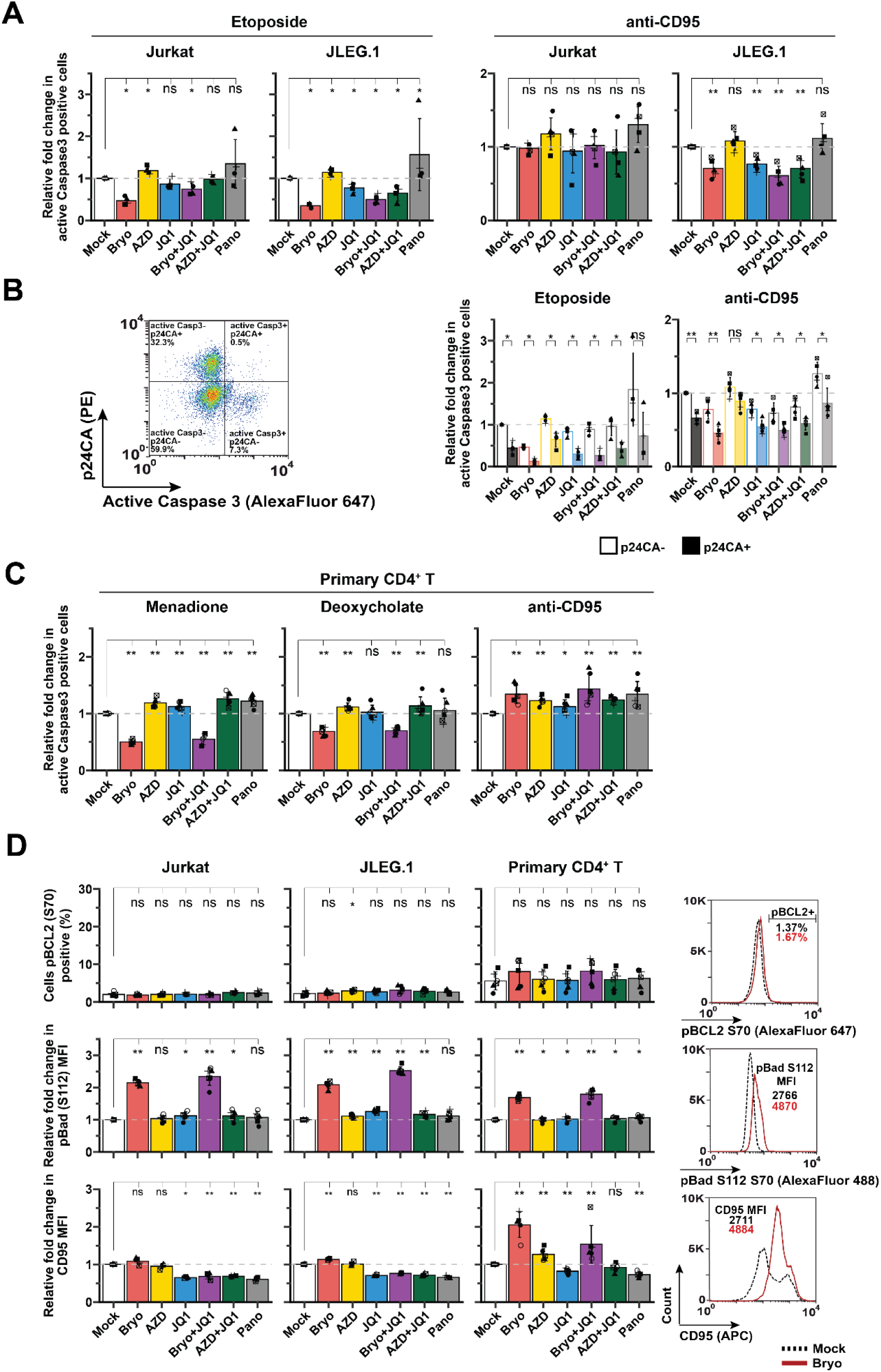
**Bryostatin-1 and the reactivating provirus both antagonise induction of apoptosis** A) Relative expression of cleaved, active Caspase 3 as a marker of apoptosis in Jurkat and JLEG.1 T-cells upon mock treatment or treatment with the indicated LRAs for 24h, followed by exposure to either etoposide (16h, left) to induce intrinsic apoptosis or an activating, anti-CD95 antibody in the presence of Protein G to induce extrinsic apoptosis (5h, right), measured by flow cytometry. Expression was normalised to the mock condition; n=4-5. B) Gating strategy used to determine the relative expression of active Caspase 3 in the p24 capsid-negative and p24 capsid-positive populations, respectively, within the same sample (left) and the relative expression of active Caspase 3 in these populations for JLEG.1 T-cells treated as in A) with the LRAs and apoptosis inducers indicated (right), measured as in A); n=4-5. C) Relative expression of active Caspase 3 in primary CD4^+^ T-cells after mock treatment or treatment with the indicated LRAs for 24h, then exposed for a further 24h to either menadione or deoxycholate to induce intrinsic apoptosis, or an activating, anti-CD95 antibody in the presence of Protein G, measured as in A); n=6. D) Expression of pBCL2 S70, pBad S112 after 1h of treatment or cell-surface CD95 expression after 24h of treatment in Jurkat, JLEG.1 and primary CD4^+^ T-cells treated with the indicated LRAs, measured by flow cytometry; n=6. Wilcoxon rank-signed test. *** p < 0.001; ** p < 0.01; * p< 0.05. Indicated is the arithmetic mean ± SD.

Experiments in primary CD4^+^ T-cells from HIV-negative donors confirmed the ability of bryostatin-1 and the bryostatin-1 and JQ1 combination to decrease intrinsically induced apoptosis levels (initiated here by treatment with menadione or deoxycholate). In contrast to a previous report which suggested that bryostatin-1 was protective against extrinsic apoptosis^37^, bryostatin-1 and the bryostatin-1 and JQ1 combination slightly but significantly increased the susceptibility of CD4^+^ T-cells to extrinsic apoptosis (Figure 5C). To understand the observed phenotypes, we analysed the phosphorylation and expression of key proteins involved in apoptotic signalling cascades in Jurkat, JLEG.1 and primary CD4^+^ T-cells. Although it has previously been reported that bryostatin-1 administration triggers the phosphorylation and associated hyperfunction of the anti-apoptotic BCL2 protein^37^, we were unable to reproduce this finding (Figure 5D). In contrast, the phosphorylation of the pro-apoptotic Bad protein at Ser112 increased upon treatment with bryostatin-1 and the bryostatin-1 and JQ1 combination across all T-cell models tested. Phosphorylation of Bad is associated with its hypofunction via cytosolic sequestration by 14-3-3^38^. Interestingly, treatment with bryostatin-1 and the bryostatin-1 and JQ1 combination significantly increased the expression of CD95 (Fas) on the surface of primary CD4^+^ T-cells, likely underpinning the observed increase in extrinsically induced apoptosis in these cells. In the same system, bryostatin-1 administration very slightly reduced expression of the anti-apoptotic BCL-xL protein, but also moderately increased levels of the anti-apoptotic MCL-1 protein (Figure S7B). In contrast, no meaningful change was observed for any combination of AZD5582 and JQ1.

Overall, this provides support to the notion that treatment with bryostatin-1 and the bryostatin-1 and JQ1 combination, but not with AZD5582 and JQ1, impedes intrinsic apoptosis pathways. Concurrently, we identified a role of the reactivating provirus in blocking both intrinsic and extrinsic apoptosis in the context of infected cells. This not only highlighted the utility of the AZD5582 and JQ1 combination strategy as more conducive to ‘shock-and-kill’, but also made clear how viral resistance to cell death signalling poses an additional hurdle that would likely have to be overcome for a maximally effective cure strategy.

### Blocks to ADCC after bryostatin-1 treatment are unrelated to ADCC co-factor modulation

Beyond the availability of cell-surface antigens and relative functioning of apoptotic pathways, an additional factor influencing ADCC-mediated killing could be downregulation of ADCC co-factors from the cell surface, including CD48, SLAMF6, PVR. However, bryostatin-1 treatment significantly upregulated the expression of CD48 in all T-cell types tested, and increased SLAMF6 expression in Jurkat and JLEG.1 cells (Figure 6A). Addition of JQ1 to Bryostatin prevented or reduced these upregulations, but did not reduce the expression level to below that of the mock-treated samples. While PVR levels were not meaningfully altered by bryostatin-1 mono-treatment or the bryostatin-1 and JQ1 combination in Jurkat cells, treatment with JQ1, AZD5582 and JQ1, and panobinostat modestly downregulated this cell-surface receptor. For JLEG.1 T-cells, PVR was more clearly downregulated in the bryostatin-1 and JQ1 condition, likely owing to HIV-1-reactivating cells showing lower cell surface PVR levels than non-reactivating cells (Figure 6B). In primary CD4^+^ T-cells, PVR levels were markedly upregulated, for both bryostatin-1 and the bryostatin-1 and JQ1 combination (Figure 6A). Bryostatin-1 also slightly upregulated the expression of HLA-E, a negative regulator of NK cell-mediated killing^39^, in Jurkat T-cells, although the combination of bryostatin-1 and JQ1 specifically led to significantly higher levels of this inhibitory factor of ADCC in primary CD4^+^ T-cells (Figure 6A). Taken together, these data indicate that low levels of ADCC observed in the bryostatin-1-containing treatments likely cannot be ascribed to ADCC co-factor modulation.

**Figure 6.**
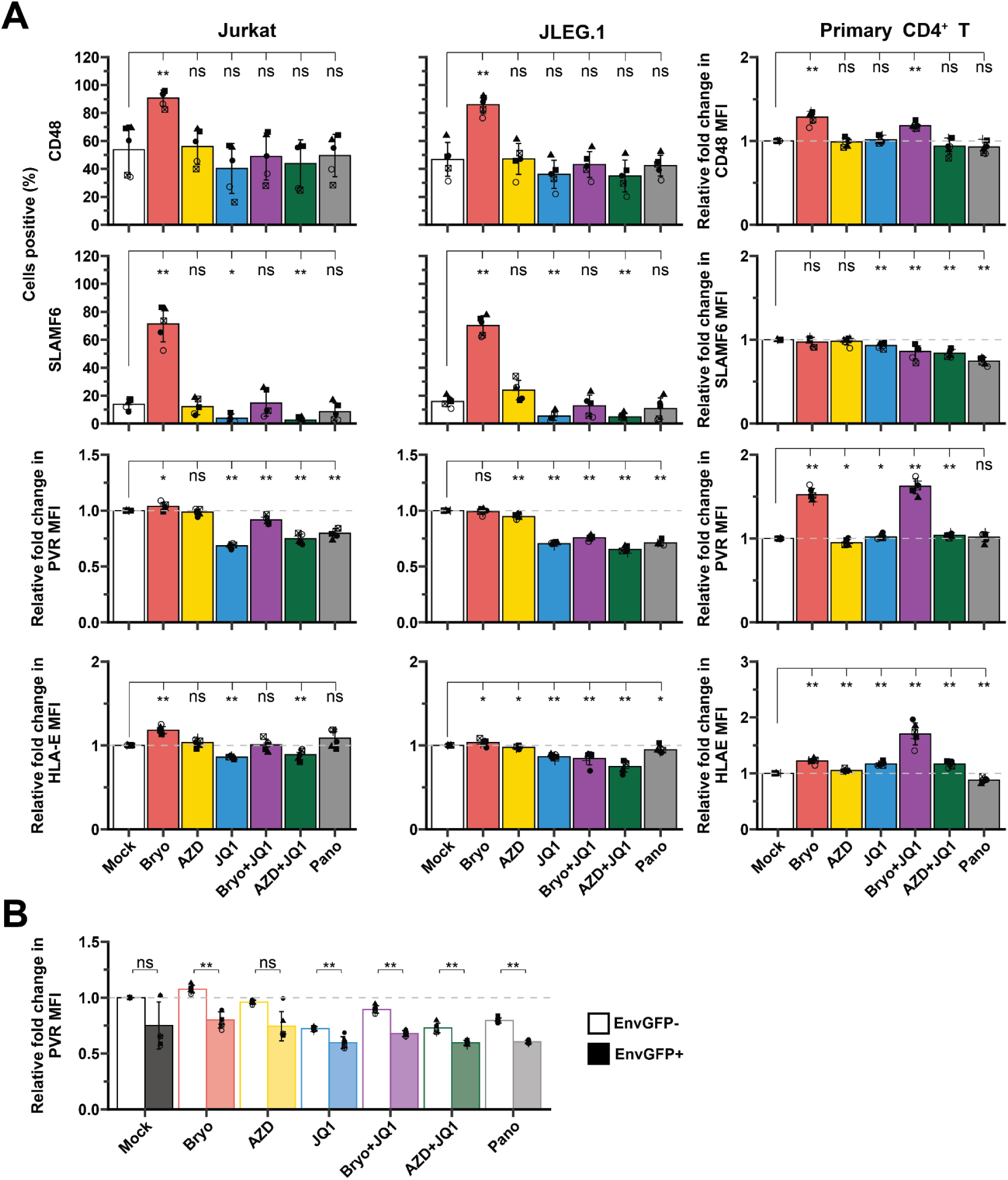
**Blocks to ADCC after bryostatin-1 treated are unrelated to ADCC co-factor modulation** A) Cell-surface expression of ADCC co-factors CD48, SLAMF6 and and PVR, as well as the inhibitory molecule HLA-E, in parental Jurkat, JLEG.1 and primary CD4^+^ T-cells treated with the indicated LRAs for 24h, measured by flow cytometry; n=6. B) Relative expression of the NK cell ligand PVR in the EnvGFP- and EnvGFP+ populations for JLEG.1 T-cells treated as in A), measured by flow cytometry. Expression was normalised to the MFI of the EnvGFP- cells in the mock condition; n=6. Wilcoxon rank-signed test. *** p < 0.001; ** p < 0.01; * p< 0.05. Indicated is the arithmetic mean ± SD.

### Knockout of the viral *nef* gene increases susceptibility of HIV-1-reactivating T-cells to apoptosis and ADCC-mediated killing

HIV-1 Nef has been described to exert both anti-apoptotic^40–42^ and pro-apoptotic^43,44^ effects, likely dependent on cellular and viral context. To test if Nef underpins the resistance of HIV-1-reactivating T-cells to apoptosis, we generated CRISPR/Cas9-mediated knockouts of the HIV-1 *nef* gene in the proviruses of the JLEG T-cell lines (Figure 7A). This allowed for an isogenic comparison of wild-type and *nef*-deficient proviruses situated at the identical integration site. As expected, *nef* knockout prevented the Nef-mediated downregulation of cell-surface MHC-I^45^ (Figure 7B) and partially rescued cell-surface CD4 levels (Figure S8A)^46^ on HIV-1-reactivatingT-cells, functionally validating *nef* knock-out. CD69 expression, however, as a marker for T-cell activation, appeared to be largely independent of Nef expression (Figure S8A). Interestingly, for most, but not all LRAs tested, loss of Nef partially reduced the levels of HIV-1 reactivation as measured by viral p24CA expression (Figure 7C, left panel), but markedly increased relative cell-surface levels of EnvGFP for all non-bryostatin-1-containing LRA treatments in two of three JLEG clones (Figure 7C right panel, Figure S8B).

**Figure 7.**
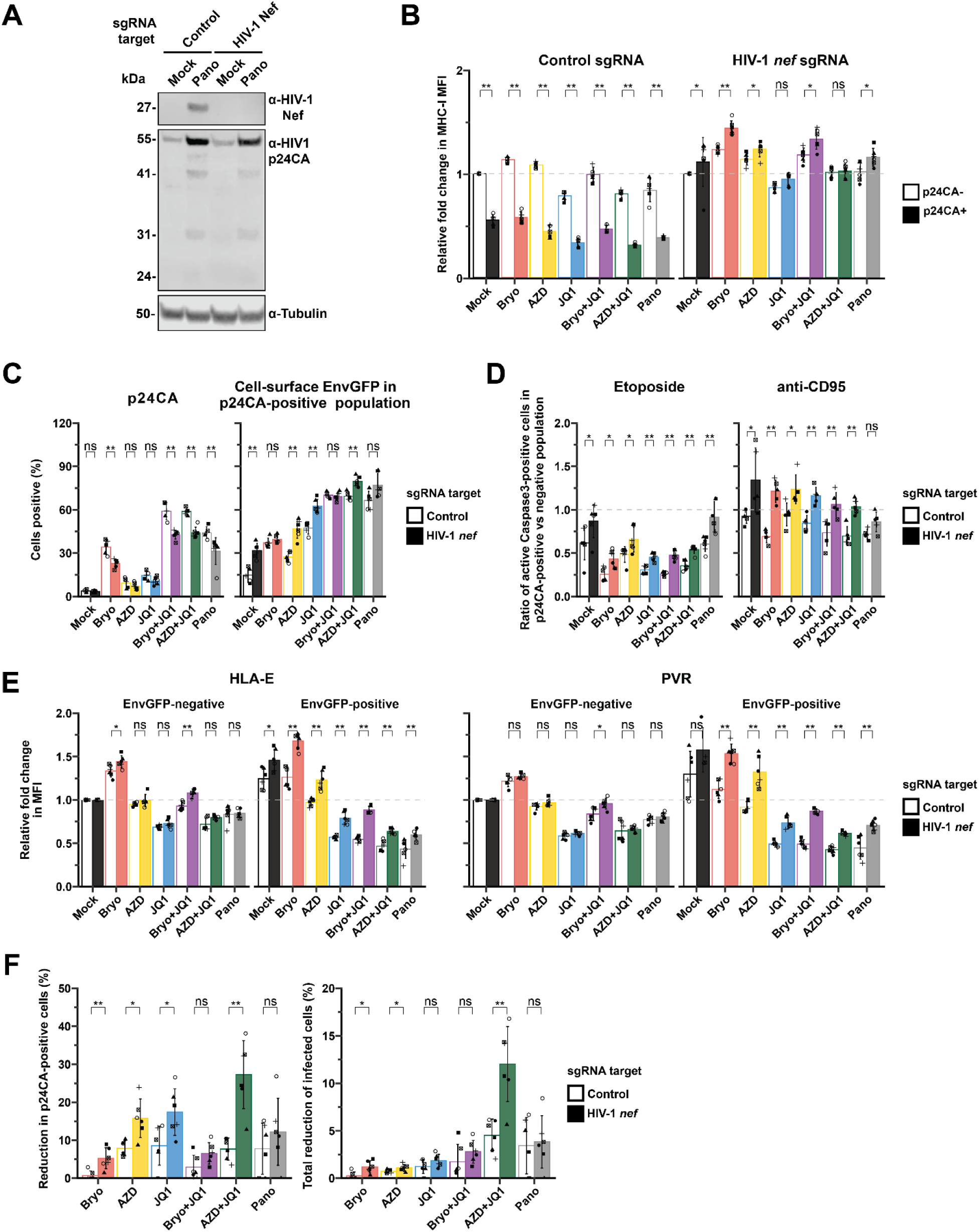
**Knockout of the viral *nef* gene increases susceptibility of HIV-1-reactivating T-cells to apoptosis and ADCC-mediated killing** A) Representative immunoblot showing the efficacy of the CRISPR/Cas9-mediated knockout of the proviral *nef* gene in JLEG.1 T-cells, either mock-treated or treated with panobinostat for 24h. The viral p55Gag protein is included as a control for treatment efficacy and Tubulin is included as a loading control. Cells were lentivirally transduced with a vector encoding Cas9 and an sgrRNA targeting *nef*, or a non-targeting, control sgRNA. B) Relative fold change in the MFI of cell-surface MHC-I in *nef* knockout and control JLEG.1 T-cells for p24-negative and p24-positive T-cells in the same sample, respectively, measured by flow cytometry. Expression was normalised to the MFI of the p24-negative cells in the mock condition; n=6. C) Expression of p24 and cell-surface EnvGFP in the p24CA-positive population of *nef* knockout and control JLEG.1 T-cells, either mock-treated or treated with the indicated LRAs for 24h, measured by flow cytometry; n=6. D) Relative expression of cleaved, active Caspase 3 as a marker of apoptosis in *nef* knockout and control JLEG.1 T-cells, mock-treated or treated for 24h with the indicated LRAs, then exposed either to etoposide (16h, left) to induce intrinsic apoptosis or an activating, anti-CD95 antibody in the presence of Protein G to induce extrinsic apoptosis (5h, right), measured by flow cytometry. Expression was normalised to the mock condition; n=5-6. E) Relative fold change in the MFI of cell-surface HLA-E (left) and PVR (right) in *nef* knockout and control JLEG.1 T-cells for EnvGFP-negative and EnvGFP-positive cells in the same sample. Expression was normalised to the MFI of the EnvGFP-negative cells in the mock condition; n=6. F) Reduction in p24CA-positive cells (left) and total reduction of infected cells for JLEG.1 T-cells treated with the indicated LRAs for 24h, followed by co-culture with PBMCs from HIV-negative donors in the presence of sera from PLWH for 4h to induce antibody-dependent cytotoxic killing; n=6 PBMC donors. Wilcoxon rank-signed test. *** p < 0.001; ** p < 0.01; * p< 0.05. Indicated is the arithmetic mean ± SD.

Loss of Nef significantly improved the susceptibility of HIV-1-reactivating T-cells to intrinsic apoptosis, though it did not completely overcome the observed resistance relative to non-reactivating cells (Figure 7D). However, in the case of extrinsically induced apoptosis, knockout of *nef* appeared to completely ablate the resistance of HIV-1-reactivating T-cells to this apoptotic signalling mode in all treatments except for panobinostat. Taken together, these results demonstrate the ability of Nef to interfere with both intrinsic and extrinsic apoptotic signalling pathways in the context of HIV-1 reactivation. In addition, we observed a pattern of higher levels of pBad S112 in HIV-1-reactivating, compared to non-reactivating T-cells (Figure S8C). Surprisingly, *nef* knockout had no effect on this phenotype, though triggering Bad phosphorylation has been described as a key mechanism by which Nef blocks intrinsic apoptosis^40^.

As knock-out of *nef* increased the susceptibility of reactivating T-cells to apoptosis and enhanced cell-surface Env levels, we investigated whether it may also bolster ADCC responses. As expected^47,48^, deletion of *nef* resulted in higher ADCC inhibitory factor HLA-E and ADCC co-factor PVR levels in HIV-1-reactivating cells (Figure 7E), where the absence of a complete rescue can likely be ascribed to Vpu in this system. For all treatments except panobinostat, *nef* knockout resulted in a significant increase in the percentage of p24CA-positive cells killed in the three JLEG cell lines in an ADCC assay (Figure 7F, left panel, Figure S8D). Crucially, the combination of AZD5582 and JQ1, in the context of *nef* knockout, resulted in the largest total reduction in infected cells for all three JLEG clones (Figure 7F, right panel, Figure S8D). This observation not only highlights the role played by Nef in the evasion of immune clearance by latently infected cells, but also outlines how the synergistically acting iIAP and iBET combination, paired with targeted Nef antagonism, could form the basis of a novel approach to HIV cure strategies that emphasises both the efficacy of latency reversal and the potency of host immune function in clearing the latent reservoir.

## DISCUSSION

Assessment of HIV latency reversal efficacy and dynamics has largely been performed in cells bearing minimal proviral reporter systems, such as the JLat model^49–51^, or proviruses often lacking both *nef* and/or *env^52–56^*. The vast majority of these cellular studies have investigated the ‘shock’ component of ‘shock-and-kill’ by assessing reporter expression following latency reversal, while not many have assessed the ‘kill’^57^. Due to the relative rarity of infected cells in peripheral blood from PLWH under ART, *ex vivo* analyses are typically restrained to quantifying changes of intra-or extracellular viral RNA levels (‘shock’) and proviral DNA levels (‘kill) at bulk level^58–60^. While these investigations are valuable, they cannot establish the percentage of cells reactivating HIV-1 transcription among all latently infected cells, and do not capture events modulating immune clearance upon latency reversal, such as Nef-mediated downregulation of MHC-I or cell-surface presentation of Env glycoproteins. The JLEG model of latency developed here is based on a full-length provirus with all viral genes expressed and functional. It more closely models authentic HIV-1 reactivation and allows tracking of both Env expression and cell-surface presentation at single-cell level. Importantly, with each cell containing a single provirus, this system allows quantification of relative efficiency of elimination upon a given treatment, something that is not achievable when using samples from PLWH. While the three T-cell clones used in this study do not represent the full spectrum of the HIV-1 reservoir heterogeneity present *in vivo*, they all shared latency reversal characteristics and responses to LRA treatment, indicating that results obtained are not specific to one specific integration site.

While a plethora of individual LRAs have been described to date^7,61^, combinations of compounds represent an opportunity to achieve synergy and target a more diverse population of proviruses whose reactivation may require different mechanisms, while potentially allowing the use of lower concentrations *in vivo*. In addition to confirming the reportedly synergistic and potent ability of the combination of the iIAP AZD5582 and iBET JQ1 to reverse latency^9^, our data establish that it is uniquely able to prevent T-cell activation, maintain full susceptibility to apoptosis induction, result in high levels of Env expression and, as a consequence induce the highest elimination rates of all treatments tested. As a bipartite strategy, ‘shock-and-kill’ necessitates that equal weighting be given to both the endeavour to reactivate transcription of latent proviruses and, in a second step, eliminate virus-infected cells. Mechanistically, knowledge regarding how treatment with LRAs, and the consequences of reactivation of proviral gene expression, affects cellular states, is limited. Furthermore, a better understanding of how the resulting perturbations modulate susceptibility to immune-mediated killing is also needed. T-cell activation, as measured by established activation markers, was clearly induced by bryostatin-1-containing treatments but not for any combination of AZD5582 and JQ1. This activation phenotype was also driven by proviral reactivation, as determined by discriminating between HIV-1-reactivating and non-reactivating cells in the same sample, and has been described to render T-cells refractory to pro-apoptotic stimuli^62–65^, thereby making them less susceptible to immune clearance. Of note was the upregulation of PD-L1 in response to the reactivating provirus, independent of the LRA applied, and to bryostatin-1 treatment directly (as evidenced in uninfected Jurkat T-cells). Of note, PD-L1 expression dampens cytotoxic CD8^+^ T-cell responses, a phenomenon best described in the context of cancer^21^, and therefore should not be boosted by an ideal LRA. To our knowledge, PD-L1 upregulation in the context of HIV-1 infection or replication as a direct consequence of intracellular viral components, rather than exogenous addition of HIV-1-derived TLR agonists ^66^ or induction of type I IFN signalling^67^, has not been described before, suggesting that this may be an unexplored route by which HIV-1-infected cells evade CTL recognition. Our observation that bryostatin-1 upregulates key T-cell activation markers in all analysed T-cell models is in line with several reports^68,69^xlink:href=" some of which however demonstrated limited proliferative responses, pointing towards incomplete T-cell activation by Bryostatin alone^68^.

Presentation of cell-surface Env as the only extracellularly accessible viral antigen is a crucial factor for ADCC responses, which hinge on the binding of anti-Env antibodies and the subsequent recruitment of effector cells^70,71^. However, we identified the HIV-1 restriction factor GBP5 to be induced by bryostatin-1 administration and in response to a reactivating provirus, though not as a direct result of AZD5582 and JQ1 combination treatment. Given its documented ability to interfere with biosynthesis and intracellular trafficking of HIV-1 Env during *de novo* infection^19,20,34^, we hypothesised high GBP5 expression in the context of latency reversal would be accompanied by reduced cell-surface Env levels. This was confirmed and *GBP5* knockout experiments established a causal relationship between GBP5 expression and reduction of cell-surface Env, demonstrating that at least a part of the bryostatin-1-imposed suppression of cell-surface Env presentation is mediated by induction of this restriction factor. It is important to note that the degree of GBP5 induction in the context of the combination of bryostatin-1 and JQ1 was insufficient to fully impede HIV-1 Env cell-surface presentation, an observation most likely explained by saturation of the restriction factor by high levels of Env expression upon reactivation by this potent combination^34^. It will be interesting to investigate by which mechanism the combination of AZD5582 and JQ1 suppresses GBP5 expression, and whether other Env-targeting ISGs, which would otherwise interfere with cell-surface Env presentation, may be equally suppressed.

Recent efforts to sensitise latently infected cells to apoptosis centre on inhibiting the anti-apoptotic BCL2 protein^72–74^. Notably, the IAP antagonist AZD5582, beyond reversing latency, can also sensitise cells to apoptosis by binding to and triggering the degradation of the anti-apoptotic proteins cIAP1 and XIAP, a phenotype described in the context of human pancreatic cancer^75^. As such, AZD5582 and related molecules have the potential to perform a dual function in HIV cure approaches. In contrast to AZD5582-containing treatments, bryostatin-1-containing treatments reduced the ability of T-cells to cleave caspase-3 upon administration of inducers of intrinsic apoptosis. While this refractoriness to apoptosis inducers is potentially linked to the initiation of T-cell activation by bryostatin-1, it was accompanied by phosphorylation of Bad at S112, a post-translational modification known to induce the hypofunction of this pro-apoptotic protein by facilitating its cytosolic sequestration via 14-3-3^38,76^. Bad phosphorylation and function depend on the MAPK/ERK and PI3K/Akt signalling pathways^77,78^. Both of these are directly involved in T-cell activation responses^79–83^, providing a likely mechanistic link between T-cell activation, Bad phosphorylation and lowered responsiveness to apoptosis inducers. Interestingly, bryostatin-1 has previously been described to block apoptosis by inducing phosphorylation and subsequent hyperfunction of the anti-apoptotic BCL2 protein at S70^37^, though this was not reproducible in our investigation. Importantly, a flow cytometric analysis differentiating LRA-treated cells according to their p24CA positivity status uncovered a propensity of HIV-1-reactivating cells to evade both intrinsic and extrinsic apoptosis, indicating a contribution of a co-induced, viral component in counteracting apoptosis induction. The combination of these bryostatin-1-induced blocks to and viral antagonism of apoptosis likely underpins the low levels of ADCC-mediated killing observed for bryostatin-1 containing conditions, though we concluded that this was likely not due to a modulation of ADCC co-factors by the LRA.

The HIV-1 viral accessory protein Nef is a multi-functional protein, perhaps best characterised for its ability to downregulate cell-surface CD4 and MHC-I levels^45,46^. It also appears to act both pro- and anti-apoptotically, depending on context^40–44^. By downregulating MHC-I, Nef allows for immune evasion by cytotoxic lymphocytes, while its downregulation of PVR ^47^ and CD4 ^84^ serves to confound humoral immunity. Our findings demonstrating Nef as a factor reducing cell-surface Env presentation are reminiscent of early work demonstrating Nef-modulated efficiency of Env-dependent syncytia formation^85^. The exact role of Nef in modulating cell-surface levels of Env remains somewhat controversial, however, with early studies suggesting that Nef expression leads to less presentation of Env^86^ and others suggesting that Vpu, rather than Nef, is responsible for limiting cell-surface Env^84^. What has become apparent is that, while cell-surface Env is required for ADCC, the efficacy of cytotoxic killing hinges on the binding of anti-Env IgG serum antibodies and this, in turn, is dependent on Env conformation and CD4-Env binding^70,87,88^. Because the CD4-bound, “open” conformation of Env exposes epitopes readily recognised by antibodies in sera from PLWH^87^, downregulation of CD4 by Nef shifts Env toward the more ADCC resistant “closed” conformation, thereby impeding ADCC^84,89^. However, while all prior work has been done in the context of acute infection, our report is the first to describe the role of Nef in modulating cell-surface Env in the context of latency reversal. In addition to diminishing antigen visibility, this work is the first to provide evidence of Nef protecting T-cells from intrinsically and extrinsically induced apoptosis in the context of latency reversal. Importantly, these Nef-mediated phenotypes culminated in Nef-mediated protection from antibody-dependent cytotoxic killing, establishing Nef as a key, yet overlooked antagonist of ‘shock-and-kill’ strategies. Indeed, it is likely that Nef-mediated antagonism of ADCC is driven by a combination of PVR downregulation, apoptosis inhibition, cell-surface Env presentation limitation and prevention of the ADCC-sensitising, “opening” CD4-Env interaction by degradation of CD4.

Together, this work advocates for a careful preclinical assessment of biological consequences, and efficacies both at ‘shock’ and ‘kill’ level, of LRA treatment before prioritisation for clinical studies. As an example, a phase I clinical trial in aviremic PLWH found that bryostatin-1 administration, while well tolerated, failed to induce a detectable increase in HIV RNA levels^90^. An analogous study investigating panobinostat demonstrated a measurable, treatment-induced upregulation of unspliced, cell-associated viral RNA but no reduction in the overall viral reservoir^10^. These results are in line with those presented in this study, where we found that bryostatin-1 had limited efficacy when administered as mono-treatment. Panobinostat, in addition, triggered robust latency reversal, but underperformed relative to the AZD5582 and JQ1 combination where ADCC-mediated elimination efficacy was concerned. In line with this, work from our group^91^ and others ^92^ has indicated that panobinostat may dysregulate normal function in immune cells. Possibly, this could also impact the susceptibility of cells to cytotoxic killing, even in the absence of a functional, ant-apoptotic Nef protein.

As a way forward, we propose synergistic latency reversal with an iIAP and iBET (AZD5582 and JQ1, or related molecules), coupled with ablation of the viral Nef protein, as a blueprint for the development of a cure strategy merging potent HIV-1 reactivation, optimal immunological visibility and full susceptibility to immune-mediated killing of infected cells. Other tested individual and combinatorial latency reversal strategies suffered from one or several insufficiencies and should therefore be deprioritised regarding *in vivo* testing. Notably, the simultaneous administration of iBETs and iIAPs *in vivo* has been reported to be well-tolerated in a small study of SIVmac251-infected macaques^93^ and induced favourable immunomodulatory effects as well as restriction of tumor growth in a mouse model of pancreatic cancer^94^, indicating good translational potential of this strategy. While some headway has been made in the development of Nef inhibitors and degraders^95–97^ and they have shown promise in potentiating immune-mediated killing of infected cells^98^, further research is needed in the context of ‘shock-and-kill’ HIV cure approaches. Combining synergistic latency reversal with Nef inhibition, therefore, represents a novel approach to HIV cure that should be investigated in the context of a more complex system, such as a small animal model of HIV-1 infection.

## MATERIALS AND METHODS

### Reagents

Bryostatin-1 (Bryo, Merck), (+)-JQ1 (JQ1, Universal Biologicals), AZD5582 (AZD, Sigma-Aldrich), panobinostat (Pano, Cayman Chemical), Phorbol 12-myristate 13-acetate (PMA, InvivoGen), Ionomycin (Io, InvivoGen), etoposide (Merck) were solved in dimethyl sulfoxide (DMSO, Stratech Scientific). Phytohaemaglutinin P (PHA) was obtained from Invivogen, while human recombinant IL-2 (IL-2) and CD3/CD28 Dynabeads^TM^ (anti-CD3/CD28 beads, Gibco) were obtained from STEMCELL and Thermo Fisher, respectively. The anti-CD3/CD28 beads were used according to the manufacturer’s recommendations (1:1 bead-to-cell ration). Efavirenz (EFV) was obtained from the NIH AIDS Reagent program. Mouse anti-Human CD95 (clone DX2) and recombinant Protein G’ were purchased from BD Biosciences and Sigma-Aldrich, respectively. Protein G was resuspended in sterile water (Merck) and stored at −20°C. Sodium deoxycholate (Merck) was dissolved in PBS (Merck) and frozen at −20°C. Menadione (Merck) was dissolved in N,N-Dimethylformamide (DMF, Sigma-Aldrich).

### Cell lines and primary cells

Cells were cultured at 37°C in a humidified incubator with 5% CO2. HEK293T cells were obtained from ATCC and cultured in Dulbecco’s modified Eagle’s medium (DMEM, Merck) supplemented with 10% heat-inactivated fetal calf serum (FCS, Merck), 100 IU/ml Penicillin/Streptomycin and 2 mM L-Glutamine (Thermofisher). Jurkat E6.1 T-cells were obtained from the NIH AIDS Reagents Program and cultured in RPMI 1640 (Merck) with 10% heat-inactivated fetal calf serum (FCS, Merck), 100 IU/ml Penicillin/Streptomycin, 2 mM L-Glutamine, 1X MEM non-essential amino acids and 1mM sodium pyruvate (both Thermo Fisher). Primary CD4^+^ T-cells were isolated from blood (buffy coats from anonymous, HIV-1-negative donors) using the EasySep™ Direct Human CD4+ T-Cell Isolation Kit (STEMCELL Technologies) according to the manufacturer’s instructions. Human PBMCs were isolated using Ficoll-Hypaque (Pancoll, PAN Biotech) centrifugation. Primary CD4^+^ T-cells were cultured in the same medium as Jurkat E6.1 T-cells, as were all JLEG T-cell lines, unless otherwise stated. JLEG T-cells were cultured in the presence of 100nM EFV to prevent viral spread in culture.

### Molecular cloning, lentiviral vectors and CRISPR/Cas9 knockout

The HIV NL4.3 V5.3 EnvGFP infectious molecular clone was created following ^12^, with the reporter protein and GS-linkers inserted 3’ of the third codon of the V5 loop domain of Env in the pNL4.3 plasmid (NIH AIDS Reagents Program). Knockouts in JLEG T-cells were obtained by lentiviral transduction using the LentiCRISPRv2 vector system (Addgene #52961). Sequences encoding a non-targeting, control sgRNA (5’-AAAAAGCTTCCGCCTGATGG-3’) and an sgRNA targeting human *GBP5* (5’-CATTACGCAACCTGTAGTTG-3’) were taken from the published sequences in the Brunello human knockout CRISPR library^99^. The *nef*-targeting sgRNA (5’-ATTGGTCTTAAAGGTACCTG-3’) was generated using the CHOPCHOP webtool^100,101^. The sequences were inserted into the LentiCRISPRv2 vector using the *BsmB*I cloning site and the insertion was validated by Sanger sequencing.

### LRA treatment

Cells were treated with bryostatin-1 (4nM) in the presence or absence of JQ1 (400nM), JQ1 (400nM) alone, AZD5582 (10nM) in the presence and absence of JQ1 (400nM), panobinostat (50nM) or a combination of PMA (0.01μg/ml) and Io (1μg/ml) for the indicated time points, unless stated otherwise. DMSO was used as a mock (or ‘vehicle’) control at a final concentration of 0.005% (v/v).

### Antibodies

All primary and secondary antibodies used in this study for both flow cytometry and immunoblotting, as well as the dilutions employed, are listed in Table S2.

### Flow cytometry

For cell-surface protein detection, cells were immunostained in FACS buffer (1% BSA, Gibco; 0.05% sodium azide, Thermofisher; in PBS) at 4°C in the dark for 30min. Cells were fixed in 4% paraformaldehyde (PFA, VWR) in PBS. For intracellular protein detection, PFA-fixed cells were either simultaneously immunostained and permeabilised in 0.1% Triton-X100 (Perkin Elmer) for 20min at RT, or first permeabilised in 90% methanol (Thermo Fisher) in dH2O for 15min on ice, followed by immunostaining for 30min at RT. Cells were incubated with appropriate secondary antibodies. A list of the detection antibodies used in this study is provided as supplementary table 1. Cell-surface EnvGFP was detected using an anti-GFP antibody conjugated to eFluor660 (clone 5F12.4, eBioscience) at a dilution of 1:300. Samples were analysed on a BD FACS Celesta (BD Biosciences) or a Cytek Aurora (Cytek) flow cytometer. Analysis of FCS files was performed using FlowJo v10 (FlowJo, LLC). JLEG.1 T-cells express very low levels of GBP5 without stimulation (typically 2-3% of cells positive) therefore upregulation of GBP5 was quantified as the change in the percentage of cells positive. In CD4^+^ T-cells, high levels of GBP5 are expressed at baseline (Figure 4B), therefore GBP5 was quantified as the change in MFI.

### Generation of HIV-1 virions and lentiviral particles

To generate authentic HIV-1 virions and pseudoviral particles, corresponding plasmids were transfected into HEK293T cells using the CalPhos™ Mammalian Transfection Kit (Takara). For generation of pseudoviral particles, the corresponding transfer plasmid was co-transfected with the packaging plasmid psPAX2 (Addgene #12260) and the VSV-G-encoding vector. 48h and 72h post-transfection, supernatant was harvested and filtered using a Stericup vacuum filtration system with a 0.22μm PES filter (Millipore). Viral particles were concentrated by ultracentrifugation over a 20% sucrose (Sigma-Aldrich) cushion at 152,982 x *g* at 4°C for 90min (SW 32 Ti swinging-bucket rotor, Beckman Coulter), resuspended in full RPMI medium and stored at −80°C. The HIV NL4.3 EnvGFP used for generation of JLEG cell lines was pseudotyped with vesicular stomatitis virus glycoprotein (VSV-G)^102^.

### Generation of latently infected Jurkat T-cell lines (JLEG)

Jurkat E6.1 T-cells were infected with VSV-G-pseudotyped HIV NL4.3 V5.3 EnvGFP at an MOI of 0.1 for six days, resulting in approximately 10% EnvGFP-positive cells as determined by flow cytometry. From that time point on, cells were maintained in medium containing 100nM EFV to prevent viral spread. Virtually no EnvGFP signal was detectable after 14 days. Cells were seeded into 96-well plates at a concentration of 0.5 cells per 100 ul per well and cultured for three weeks. The resulting single-cell clones were screened for the presence of a single integrated provirus per cell using nested *Alu-gag*, R-U5 PCR^103^. Positive clones were expanded and termed Jurkat Latent EnvGFP (JLEG) cells.

### Analysis of JLEG integration sites and proviral sequences

To determine the viral integration site and full proviral sequence in three provirus-positive clones used in this study, total genomic DNA was extracted and DNAseq libraries were prepared using the KAPA HyperPrep kit (Roche). To target library fragments comprising viral DNA and host-virus junctions, probe-based enrichment was performed using the KAPA HyperCap kit (v3.2, Roche), along with custom biotinylated probes to the viral DNA sequence. Libraries were quantified using a Qubit 3 fluorometer (Thermofisher) and size distributions of library fragments were validated using an Agilent TapeStation automated capillary electrophoresis machine. Libraries were multiplexed and sequenced on an Illumina Miseq platform. Dual indexed, paired end (2×150bp) reads were generated with a targeted read-depth of at least two million reads per original sample. Raw paired-end sequencing reads were aligned to the Homo sapiens reference genome (GRCh38) using Nvidia Clara Parabricks (v4.0.1). The Parabricks fq2bam workflow was used to accelerate the alignment process on GPUs, generating sorted and indexed BAM files. Following alignment, viral integration events were detected using the GRIDSS VIRUSBreakend pipeline^104,105^. This tool employs single breakend variant calling to identify viral DNA presence and genomic integration sites with high sensitivity. The resulting VCF files were then filtered using bcftools to isolate unique integration sites^106^. Strict filtering criteria were applied to retain only variants matching the 5’ and 3’ Long Terminal Repeat (LTR) boundaries of the HXB2 reference genome (NC_001802.1), ensuring the capture of integration junctions regardless of viral orientation. For this step, the HXB2 reference genome (NC_001802.1) was utilised rather than the experimental NL4.3 strain, as it is the annotated standard available in the VIRUSBreakend reference database.

### Immunoblotting

Cells were lysed using either M-PER™ Mammalian Extraction Reagent or RIPA Lysis and Extraction Buffer (both Thermofisher), supplemented with 1X Halt™ Protease Inhibitor Cocktail and EDTA and 1X Halt™ Phosphatase Inhibitor Cocktail (all ThermoFisher) for 30min at 4°C. The insoluble fraction was pelleted at 20,000*g* for 10min at 4°C and the supernatant mixed with 4X SDS lysis buffer (0.625M Tris/HCl pH6.8, Sigma-Aldrich; 8% SDS, Cytiva; 400mM β-mercaptoethanol, Universal Biologicals; 40% glycerol, Thermo Fisher; 0.08% bromophenol blue, VWR) to a final concentration of 1X. Samples were heated at 95°C for 10min and stored at −20°C. Immunoblotting and subsequent quantification of protein abundance was performed as previously described ^91^.

### Infection and lentiviral transduction

Infection of Jurkat E6.1 T-cells and lentiviral transduction of JLEG T-cells were performed by spinoculation. In brief, cells were mixed with concentrated lentiviral stocks and centrifuged at 1,000 x *g*, 37°C for 90min. Treatment with puromycin (Sigma-Aldrich) at 1μg/ml started three days post infection for ten days.

### Apoptosis assays

In T-cell lines, intrinsic and extrinsic apoptosis was induced by cultivation of cells in medium containing 20μM etoposide for 16h and 6.25μg/ml anti-CD95 (clone DX2) with 3.25 μg/ml recombinant Protein G’ for 5h, respectively. Apoptosis was induced after treatment with indicated LRAs for 24h. For primary CD4^+^ T-cells, apoptosis was induced by cultivation in medium containing 15μM menadione or 600μM deoxycholate (intrinsic apoptosis), or 6.25μg/ml anti-CD95 with 6μg/ml recombinant Protein G’ (extrinsic apoptosis) for 24 hours. Cleaved, active Caspase 3 as an indicator of apoptosis was quantified flow cytometrically by anti-active Caspase 3 immunostaining (clone C92-605.rMAb, BD Biosciences). During apoptosis induction, cells were cultured in the absence of sodium pyruvate and L-Glutamate as these may act as reducing agents.

### Preparation and sequencing of RNAseq libraries

Two million JLEG.1 T-cells and cells derived from an uninfected clone were mock-treated, or treated with bryostatin-1 (4nM), JQ1 (400nM) or a combination of bryostatin-1 and JQ1 for 24h. Cell lysates were inactivated in 300μl TRI Reagent (Zymo Research) for 10min at RT. RNA was extracted using Direct-zol RNA Miniprep kit (Zymo Research) and quantified using the Qubit™ RNA High Sensitivity kit and the Qubit 4 Fluorometer (Thermo Fisher). 400ng of RNA per sample was used to generate total RNA-sequencing (RNAseq) libraries using the Stranded Total RNA Prep with Ribo Zero Plus library prep kit (Illumina). The libraries were quality-controlled using a 4200 TapeStation device and the D1000 DNA ScreenTape kit (Agilent) and sequenced to a depth of 50 million reads per library on a NovaSeq 6000 (Illumina) as 75 nucleotide paired-end reads.

### ADCC assays

JLEG T-cells (target cells) were treated with the respective LRAs or LRA combinations for 24h, resuspended in serum-free RPMI containing a 1:1000 dilution of 1mM CellTracker™ Deep Red (Thermo Fisher) and incubated at 37°C for 15min. After washing with PBS to remove remnants of LRAs in the supernatant, stained cells were seeded at a concentration of 10^6^ cells/ml in 100μl in a 96-well plate. After addition of pooled anti-HIV sera (#3957, NIH AIDS Reagent Program, final dilution 1:100), freshly isolated PBMCs from anonymous, HIV-uninfected donors (effector cells) were added to target cells to a final effector:target cell ratio of 1.5:1. Cocultures without HIV serum served as a reference. Cocultures were centrifuged at 300g for 1min to maximise contact between effector and target cells and incubated for 4h at 37°C, 5% CO2. Following fixation, cells were immunostained for intracellular HIV-1 p24 expression (clone KC57, Beckman Coulter). The measures of ADCC efficiency were calculated as follows::

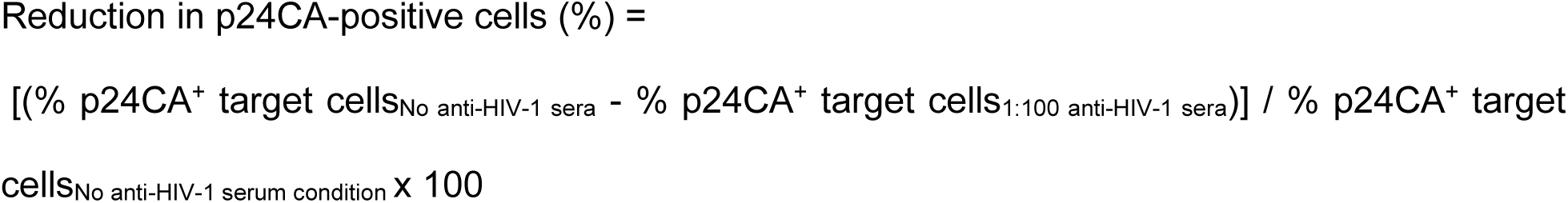

The total reduction in infected cells was calculated as follows:

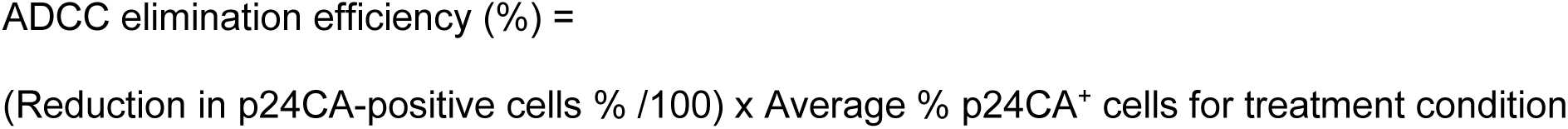

### Data analysis

FASTQ files from the sequencing output were aligned to GRCh38 v109 (Ensembl) using STAR^107^. Gene counts were assigned using Rsubread^108^ and downstream analysis was performed using DESeq2^109^ within the R programming language, v4^110^. The code is available at https://github.com/GoffinetLab/HIV_Synergistic-Latency-Reversal-study and raw sequencing data will be made available on the NCBI GEO database.

### Statistical analysis

All data was processed using R v4^110^ and graphics were produced using the ggplot2^111^ and pheatmap^112^ packages. Statistical analysis was performed using the rstatix package^113^ and Bliss-Independence testing for synergy between bryostatin-1 and JQ1 was performed using the BIGL package^114^. Bliss-Independence testing of synergy between AZD5582 and JQ1 was performed by first subtracting the value for a given parameter observed in the mock treatment from the value observed for each LRA treatment respectively. Thereafter, the expected combination effect was determined with the corrected values using the formula Effect_(expected, AZD5582+JQ1)_ = E_(observed, AZD5582)_ + E_(observed, JQ1)_ - E_(observed, AZD5582)_E_(observed, JQ1)_. Bliss excess was then calculated using the formula: Bliss excess = Effect_(observed, AZD5582+JQ1)_ - Effect_(expected, AZD5582+JQ1)_, with positive and negative values indicating synergy and antagonism respectively. Statistical significance was then determined by comparing the values for Effect_(observed, AZD5582+JQ1)_ and Effect_(expected, AZD5582+JQ1)_ using a Wilcoxon rank-signed test. Code used is available at https://github.com/GoffinetLab/HIV_Synergistic-

### Latency-Reversal-study

#### Ethics

Withdrawal of blood samples from anonymous, HIV-negative human donors and cell isolation were conducted with approval of the local ethics committees (Ethical review committee of Charité Berlin, vote EA4/167/19; NHSBT, acc. M252).

## RESOURCE AVAILABILITY

*Lead contact.* Further information and requests for resources should be directed to and will be fulfilled by the lead contact, Christine Goffinet.

*Materials availability*. Cell lines will be provided on reasonable request. *Data and code availability*. Code used in this study is available at https://github.com/GoffinetLab/HIV_Synergistic-Latency-Reversal-study.

Raw sequencing data will be made available on the NCBI GEO database.

## AUTHOR CONTRIBUTIONS

DP generated the majority of the data. STH conducted the integration site analysis of the proviruses of the three JLEG clones. BA generated the recombinant NL4.3 Env.GFP_OPT_ molecular clone. JJ and DP provided technical assistance. OC provided essential reagents. CG supervised the work. DP and CG wrote the manuscript. All authors read, edited, and approved the manuscript.

## Supporting information

Postmus et al., Supplementary Figures

## ACKNOWLEDGEMENTS

We thank the anonymous blood donors for providing their blood for research. We thank Valerie Bosch and the NIH AIDS Research & Reference Reagent Program for providing essential reagents. We thank the Genomics Platform of the Berlin Institute of Health for next-generation sequencing. This work was supported by funding to C.G. by Berlin Institute of Health (BIH); by Hector Foundation, project M2101; by DFG priority program 1923 “Innate Sensing and Restriction of Retroviruses,” grant GO2153/4; by DZIF, TTU HIV, grant 04.820 to C.G., by a Professorship Award (APR9/1017) of the Academy of Medical Sciences awarded to C.G. O.C. is an Einstein Independent Researcher (ESR-2023-770) funded by the Einstein Foundation Berlin. Additionally, this project was supported by funding to D.P. by award of the Jean Clayton Fund 2025, Liverpool School of Tropical Medicine, and the Rosetrees Seedcorn Award 2025 for the project “Targeting HIV Accessory Proteins Toward Therapeutic Intervention and Cure (TAcTIC)”.

