## Supplementary material for "Synergistic latency reversal by the IAP antagonist AZD5582 and BET inhibitor JQ1, combined with Nef ablation, facilitates immune-mediated elimination of latently HIV-1-infected T-cells": Postmus et al., Supplementary Figures

Figure S1 Postmus *et al.*, 2026

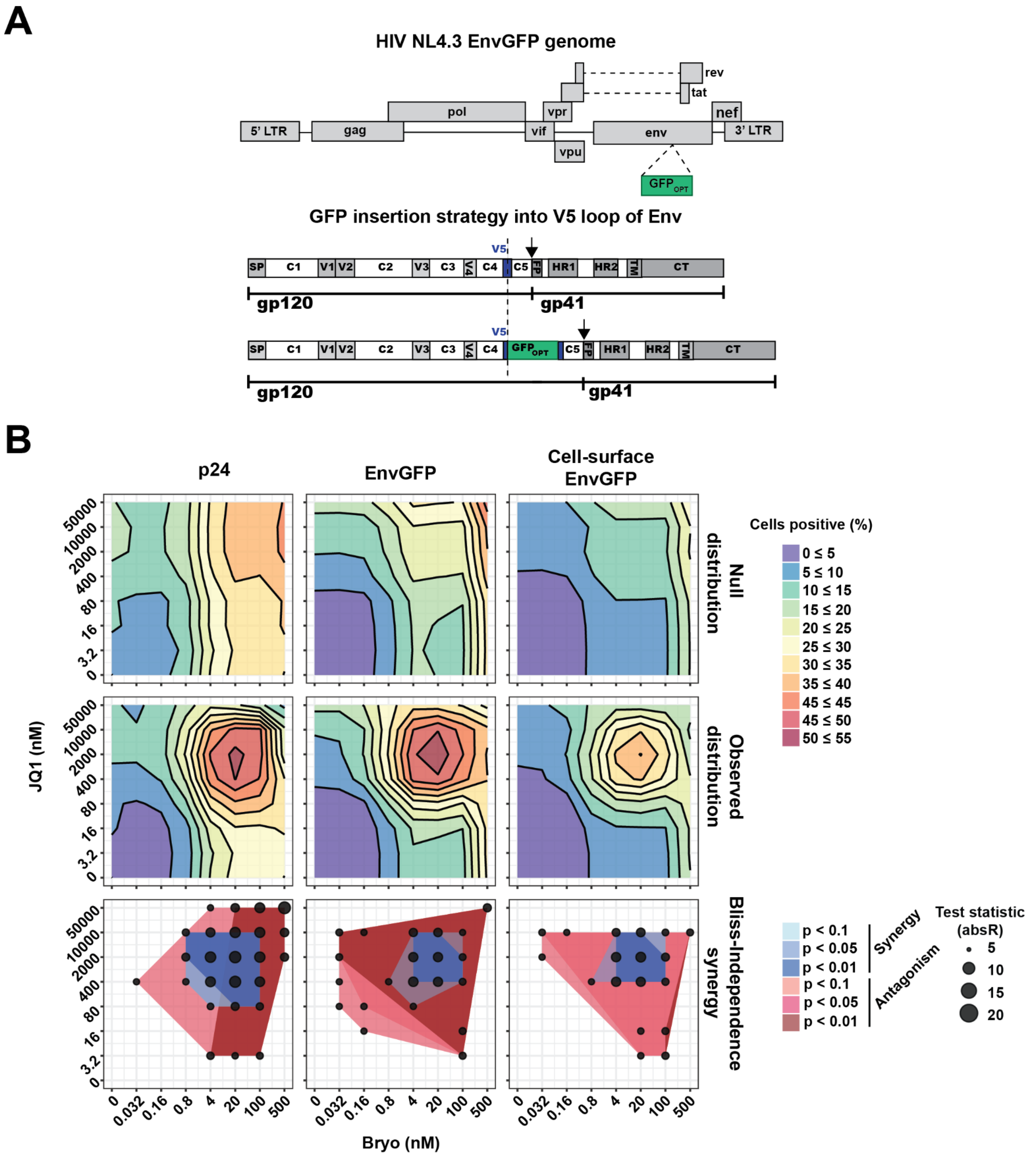


**Figure S1. Bryostatin-1 and JQ1 co-treatment synergistically reverses HIV-1 latency in JLEG.1 T-cells**

A) Schematic representation of $GFP_{OPT}$ insertion into the V5 loop domain of the full-length HIV-1 NL4.3 molecular clone to create the HIV NL4.3 EnvGFP reporter virus used to generate the latent Jurkat T-cell lines (JLEG). SP: signal peptide; V1-5: variable loops 1-5; C1-5: constant domains 1-5; FP: fusion protein; HR1-2: heptad repeats 1-2; TM: transmembrane domain; CT: cytoplasmic tail; $GFP_{OPT}$: GFP with optimised folding; dashed line: insertion site of $GFP_{OPT}$; arrow: furin cleavage site.

B) JLEG.1 T-cells were treated with combinations of bryostatin-1 and JQ1 for 24h and the expression of intracellular p24CA, total EnvGFP and cell-surface EnvGFP were quantified by flow cytometry. Indicated is the percentage of cells positive for each respective readout for the null distribution (theoretical additive distribution, extrapolated from the monotherapeutic titrations - increasing concentration of bryostatin-1 in the absence of JQ1, or vice versa, top row) or the observed distribution (middle row). Indicated is also the test statistic for synergy or antagonism, along with the corresponding p-value, for every treatment concentration combination for each respective readout using the Bliss Independence model, calculated using the BIGL R package.

Figure S2 Postmus *et al.*, 2026

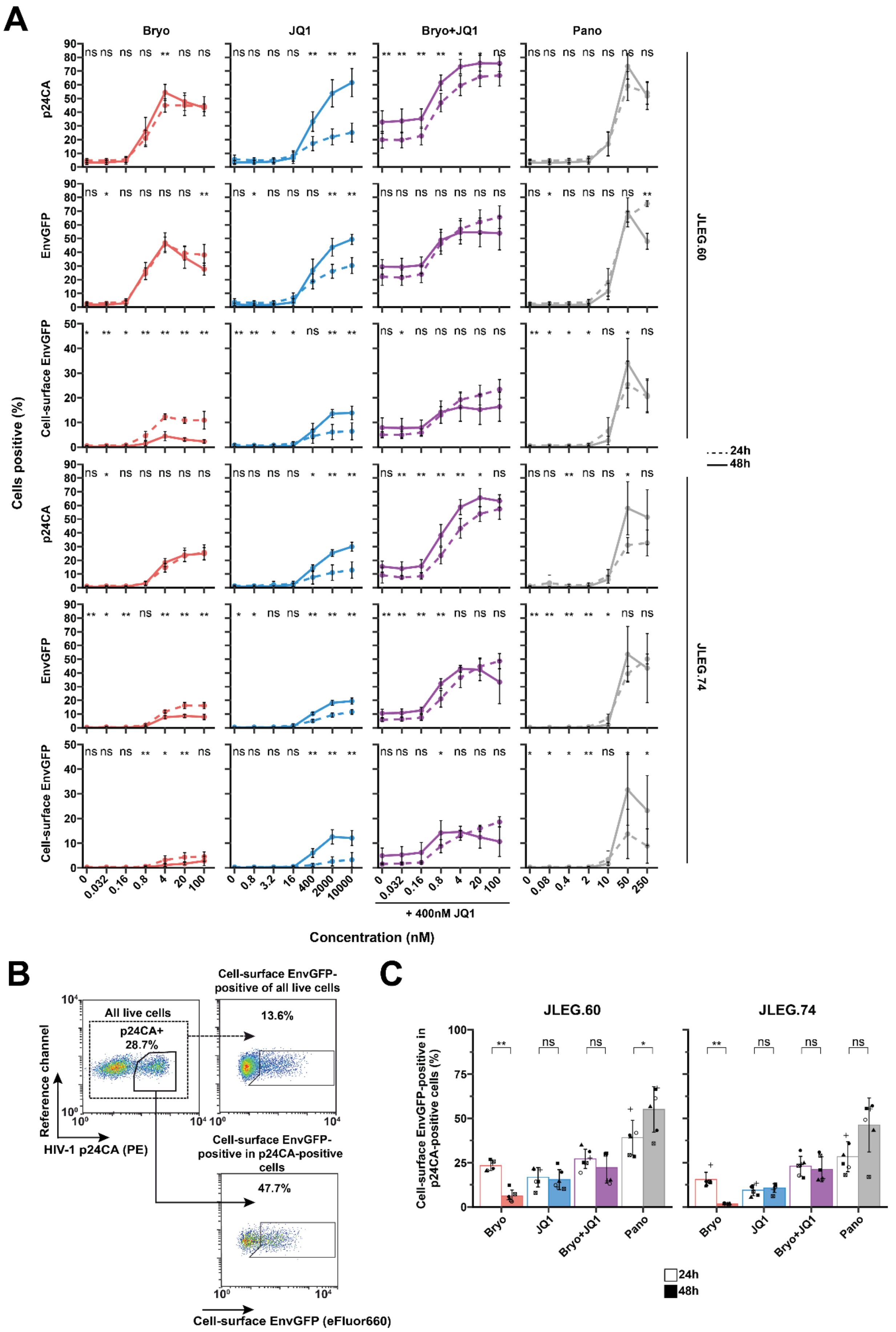

**Figure S2. Bryostatin-1 treatment decreases cell-surface EnvGFP levels in JLEG.60 and JLEG.74 T-cell clones**

A) Titration of bryostatin-1, JQ1, bryostatin-1 in the presence of 400nM JQ1, and panobinostat on JLEG.60 (top) and JLEG.74 (bottom) T-cells, measuring intracellular p24 capsid, total EnvGFP and cell-surface levels of EnvGFP at 24h and 48h post-reactivation by flow cytometry; n=6.

B) Representative flow cytometry gating strategy employed to determine the percentage of cell-surface EnvGFP-positive cells within all live singlets and specifically within the p24CA-positive subpopulation.

C) Quantification of cell-surface EnvGFP-positive cells within the p24 capsid-positive population in JLEG.60 (left) and JLEG.74 (right) T-cells treated for 24h and 48h with bryostatin-1 (4nM), JQ1 (400nM), a combination of bryostatin-1 and JQ1, or panobinostat (50nM), measured by flow cytometry; n=6 (right)..

Wilcoxon rank-signed test. *** $p < 0.001$; ** $p < 0.01$; * $p< 0.05$. Indicated is the arithmetic mean ± SD.

Figure S3 Postmus *et al.*, 2026

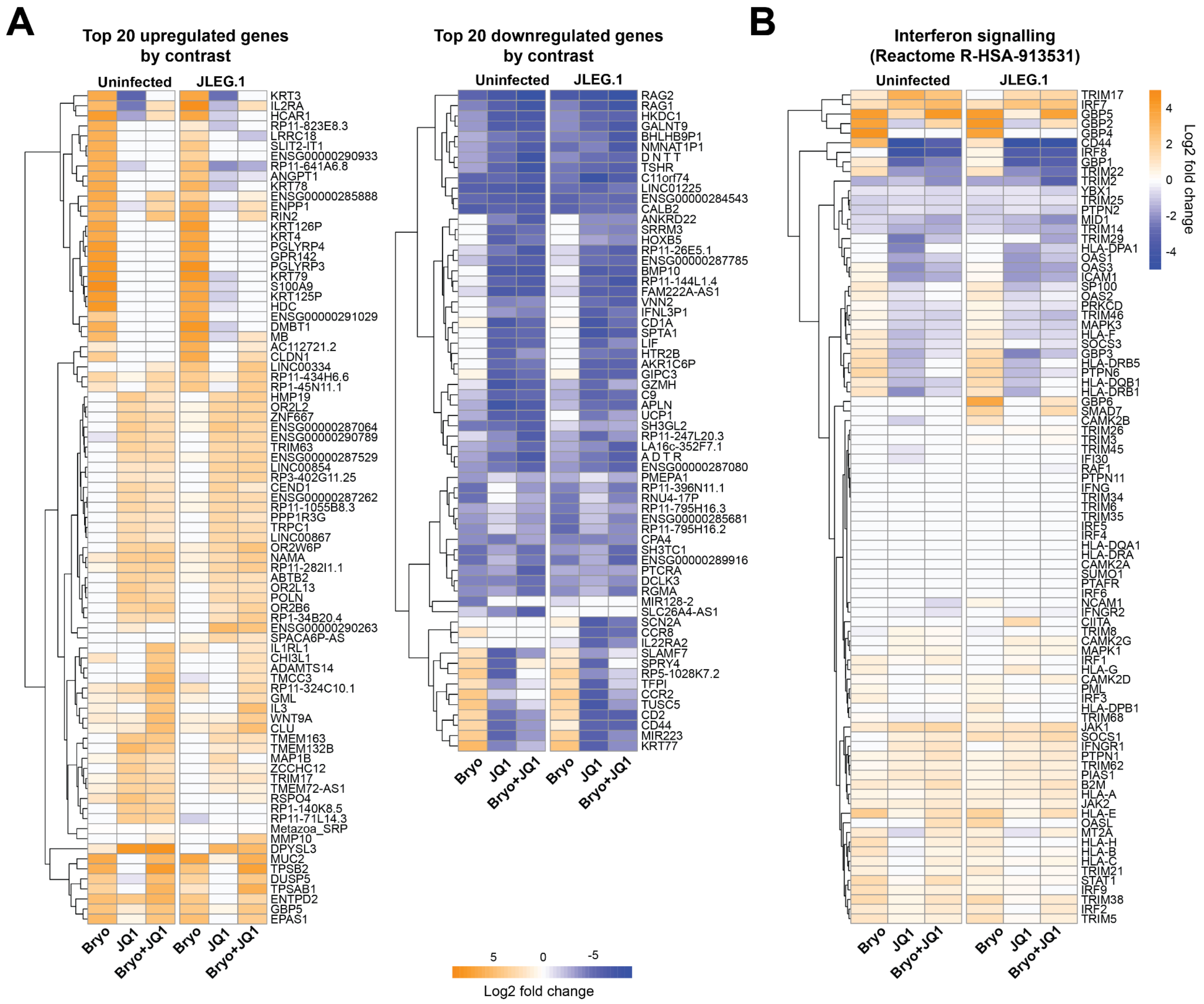


**Figure S3. Top up- and down-regulated genes and regulation of interferon-stimulated genes**

A) RNAseq analysis of JLEG.1 T-cells and an uninfected clone, treated as in Fig. 2, showing the log2 fold change in expression for the top 20 up- (left) and downregulated (genes) for each indicated treatment compared to the mock condition, selected for each contrast and clone combination.

B) Log2 fold change in expression of genes in the interferon signalling pathway (Reactome R-HSA-913531) for each treatment relative to the mock condition in the JLEG.1 clone and an uninfected clone.

Figure S4 Postmus *et al.*, 2026

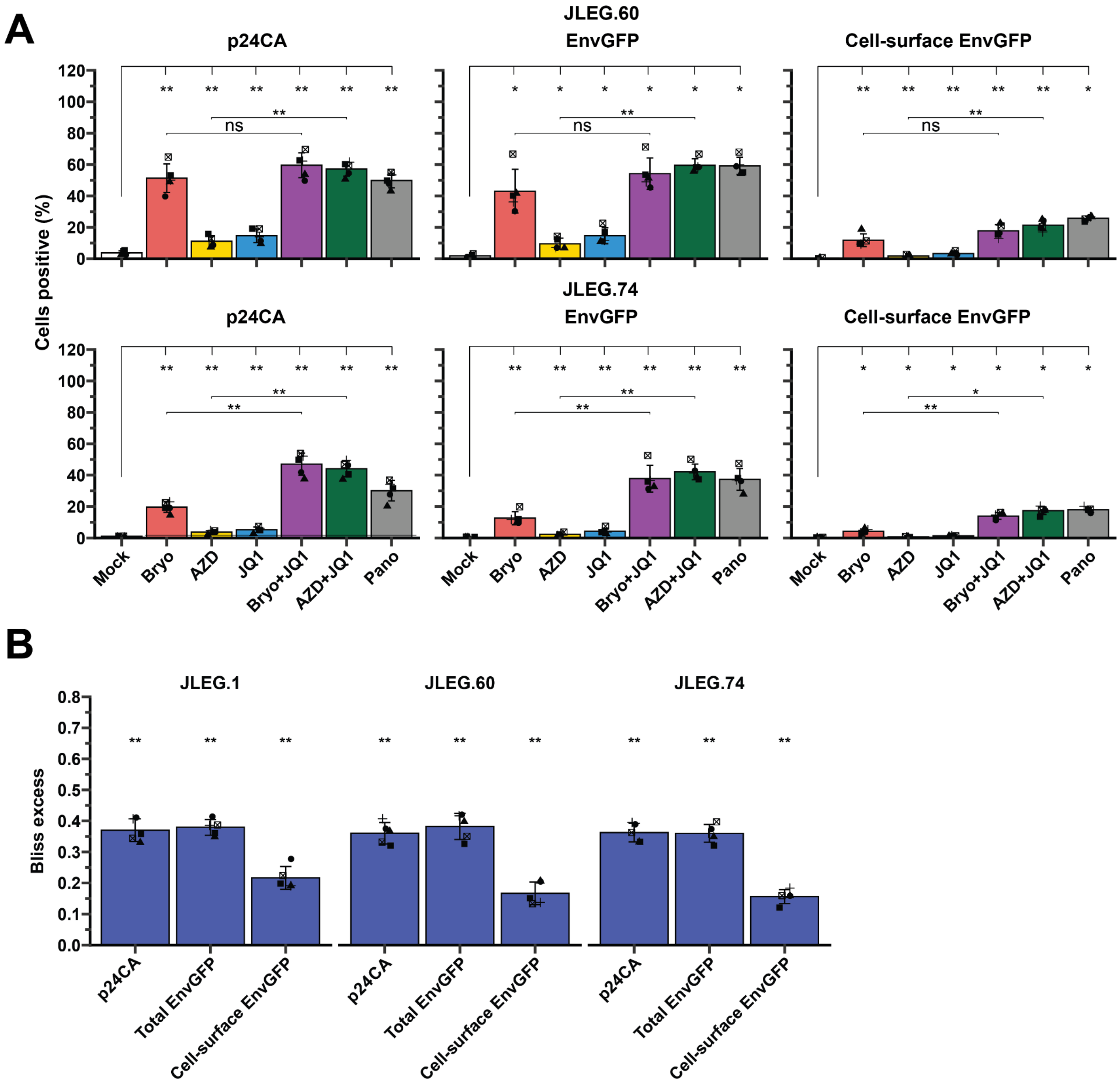


**Figure S4. Co-treatment with AZD5582 and JQ1 synergistically reverses HIV-1 latency in JLEG.60 and JLEG.74 T-cells.**

A) Expression of HIV-1 p24CA (left panel), total EnvGFP (middle panel) and cell-surface EnvGFP (right panel) in JLEG.60 (top row) and JLEG.74 (bottom row) T-cells treated as indicated for 24h, quantified by flow cytometry; n=5.

B) Bliss independence analysis of expression levels of HIV-1 p24CA, total EnvGFP and cell-surface EnvGFP in JLEG.1, JLEG.60 and JLEG.74 T-cells treated with AZD5582, JQ1, or a combination of AZD5582 and JQ1 for 24h, measured by flow cytometry. The underlying data points used are presented in A). Indicated is the Bliss excess value for each parameter and each clone respectively. Tests for statistical significance are performed by comparing the observed response for the combination treatment to the projected additive response under the Bliss independence model.

Wilcoxon rank-signed test. *** $p < 0.001$; ** $p < 0.01$; * $p < 0.05$. Indicated is the arithmetic mean ± SD.

Figure S5 Postmus *et al*., 2026

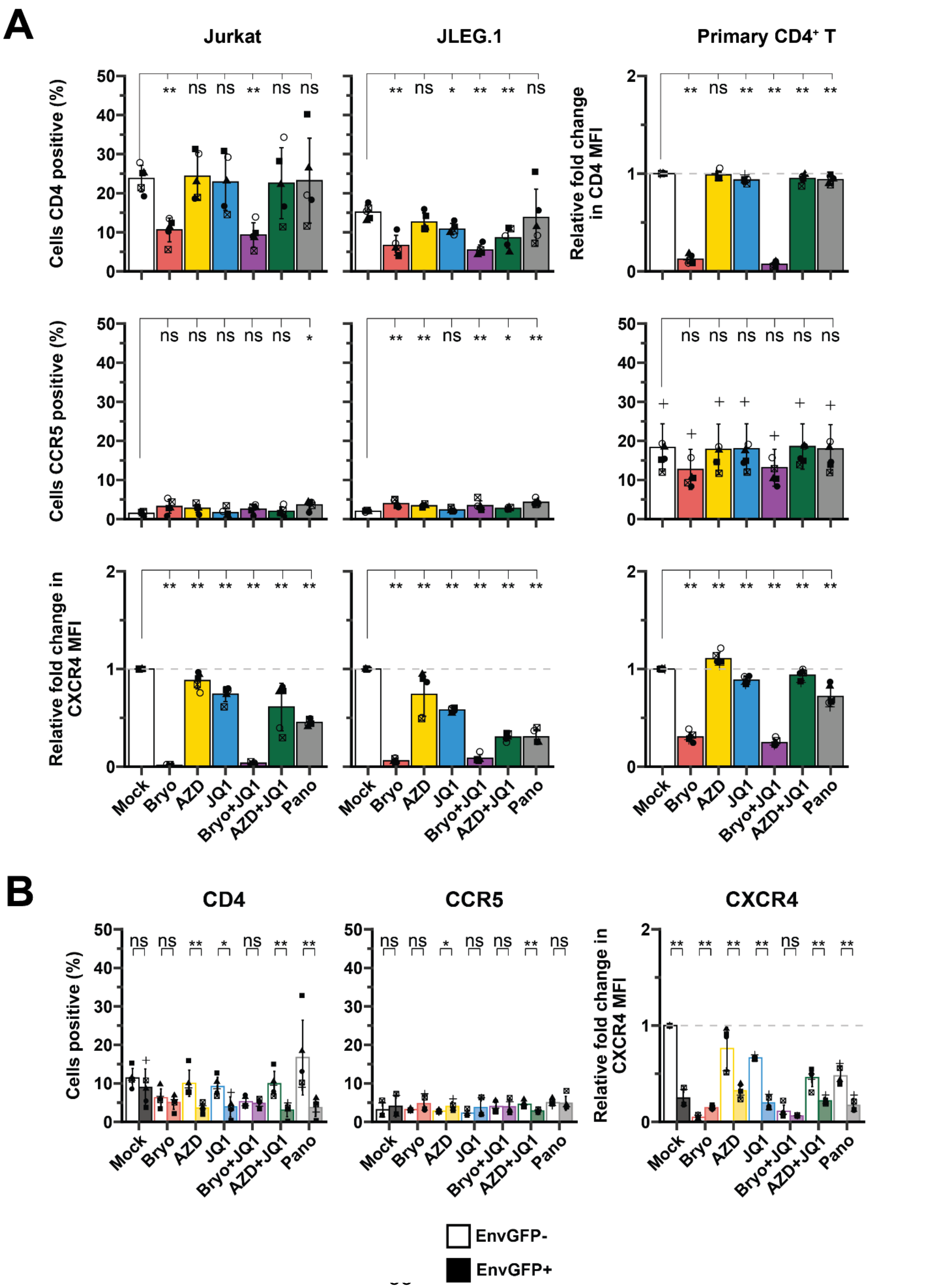

**Figure S5. Bryostatin-1 treatment downregulates cell-surface expression of CD4 and CXCR4**

A) Cell-surface expression of CD4, CCR5 and CXCR4 in Jurkat, JLEG.1 and primary CD4$^+$ T-cells treated for 24h with the indicated LRAs for 24h, measured by flow cytometry; n=5-6.

B) Cell-surface expression of CD4, CCR5 and CXCR4 in JLEG.1 T-cells for EnvGFP negative and EnvGFP positive cells in the same sample, measured by flow cytometry; n=5.

Wilcoxon rank-signed test. *** $p < 0.001$; ** $p < 0.01$; * $p< 0.05$. Indicated is thearithmetic mean ± SD.

Figure S6 Postmus *et al.*, 2026

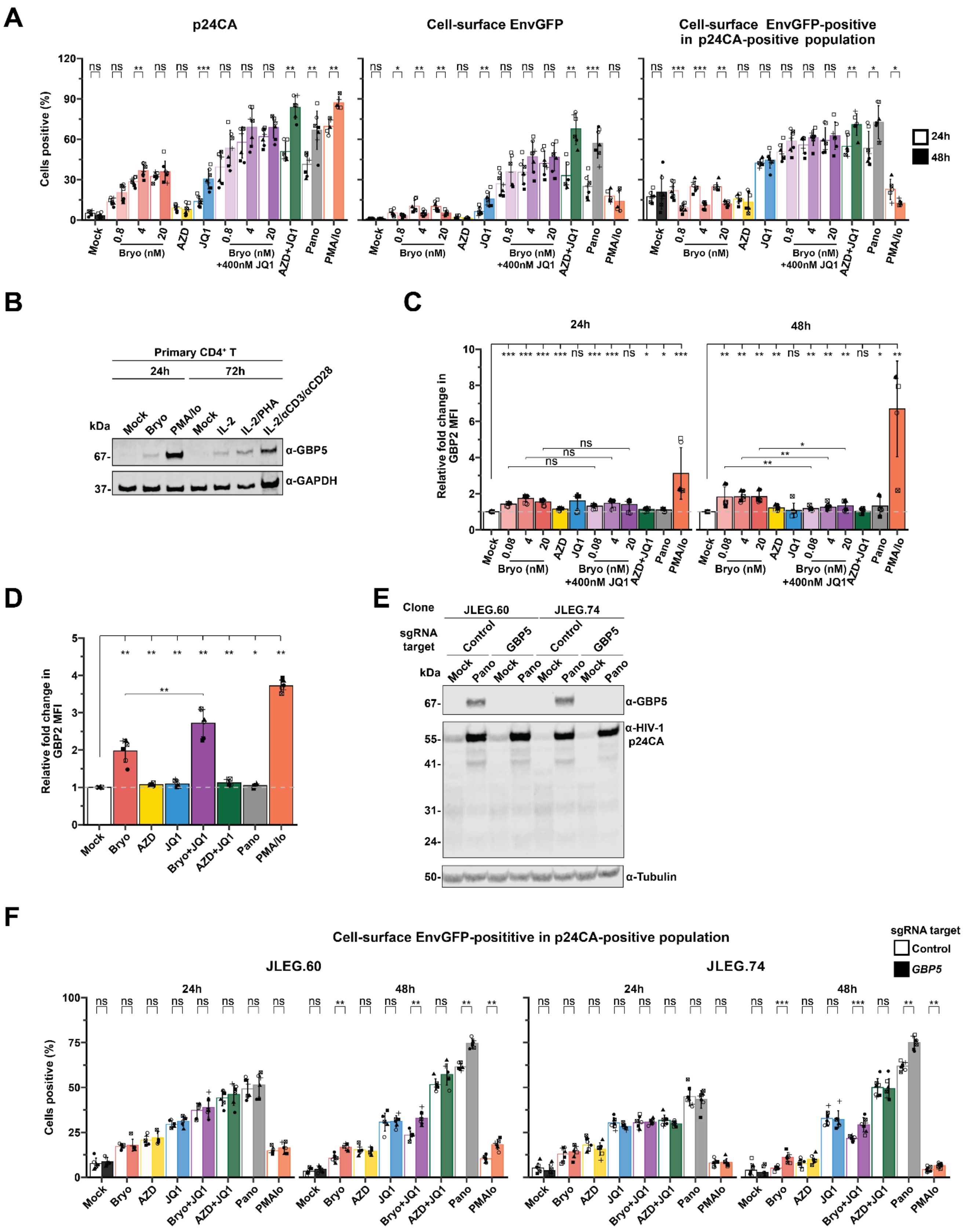

**Figure S6. Bryostatin treatment-induced GBP expression modulates cell-surface EnvGFP expression**

A) Expression of HIV-1 p24 capsid (left panel), cell-surface EnvGFP (middle panel) and cell-surface EnvGFP within the p24 capsid-positive population (right panel) in JLEG.1 T-cells after mock treatment or treatment with the indicated LRAs, including varying, indicated concentrations of bryostatin-1, for 24h and 48h, quantified by flow cytometry; n=6-7.

B) Expression of GBP5 in primary $CD4^+$ T-cells from an HIV-negative donor either mock-treated or treated with bryostatin-1 or PMA/Io for 24h, or either mock-treated or treated with IL-2 (10ng/ml), a combination of IL-2 and PHA (1ug/ml) or anti-CD3/anti-CD28 beads for 72h, assessed by immunoblotting. GAPDH is included as a loading control.

C) Expression of GBP2 in JLEG.1 cells treated as in A), quantified by flow cytometry; n=5-7.

D) Expression of GBP2 in primary $CD4^+$ T-cells treated for 24h with the indicated LRAs, quantified by flow cytometry; n=6.

E) Representative immunoblot showing the efficacy of the CRISPR/Cas9-mediated knockout of the *GBP5* gene in JLEG.60 and JLEG.74 T-cells, either mock-treated or treated with panobinostat as a known inducer of GBP5 for 24h. The viral Gag proteins (detected by anti-p24CA antibody) are included as a control for reactivation efficacy. Tubulin is included as a loading control.

F) Expression of cell-surface EnvGFP in the HIV-1 p24 capsid-positive population for *GBP5* knockout or control JLEG.60 and JLEG.74 T-cells after mock treatment or treatment with the indicated LRAs for 24h and 48h.

Wilcoxon rank-signed test. *** $p < 0.001$; ** $p < 0.01$; * $p < 0.05$. Indicated is the arithmetic mean ± SD.

Figure S7 Postmus *et al.*, 2026

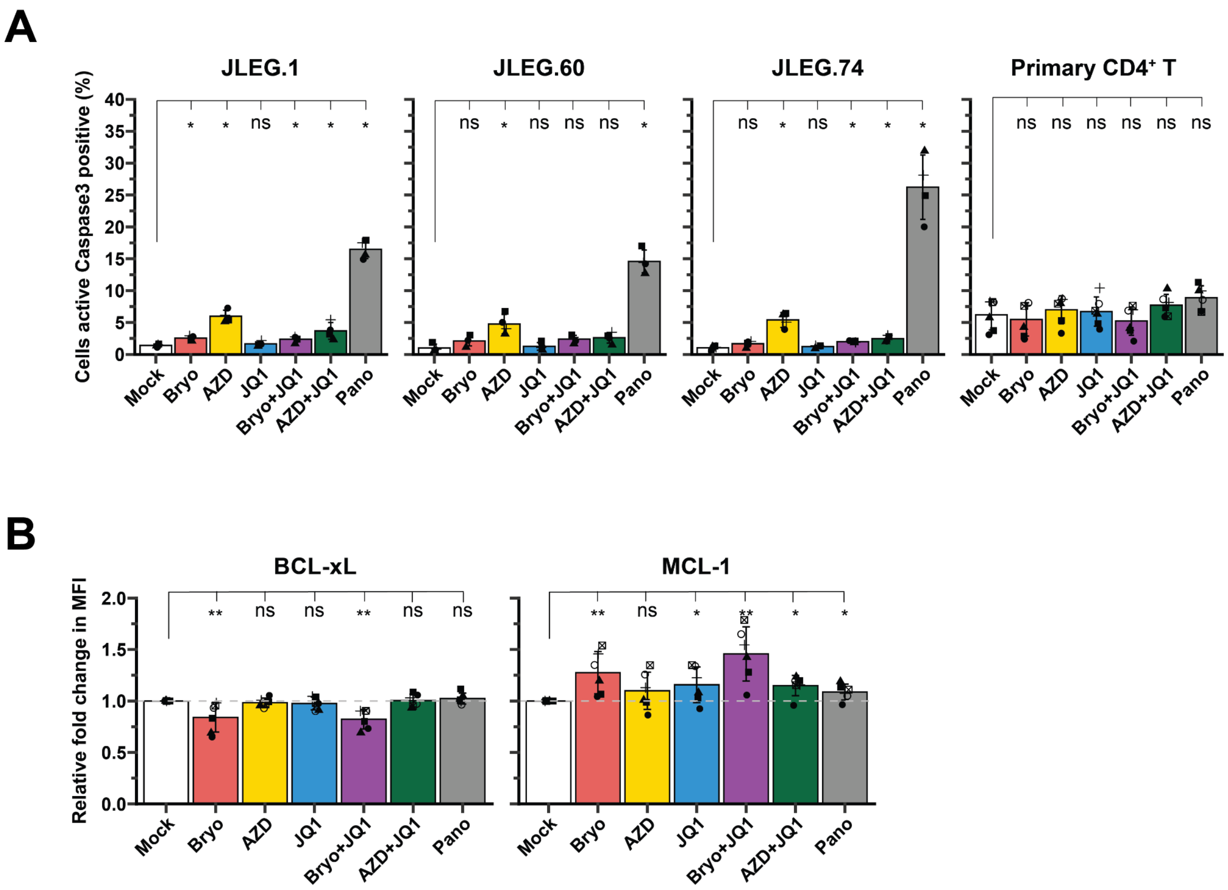


**Figure S7. Panobinostat treatment induces high levels of toxicity in JLEG cells and bryostatin-1 triggers upregulation of the anti-apoptotic MCL-1 protein in primary CD4⁺ T-cells**

A) Expression of active Caspase 3 as a measure of treatment cytotoxicity in JLEG.1, JLEG.60, JLEG.74 and primary CD4⁺ T-cells treated with the indicated LRAs for 24h; n=4.

B) Expression of BCL-xL and MCL-1 in primary CD4⁺ T-cells either mock-treated or treated with the indicated LRAs for 24h, measured by flow cytometry; n=6.

Wilcoxon rank-signed test. *** $p < 0.001$; ** $p < 0.01$; * $p < 0.05$. Indicated is the arithmetic mean ± SD.

Figure S8 Postmus *et al.*, 2026

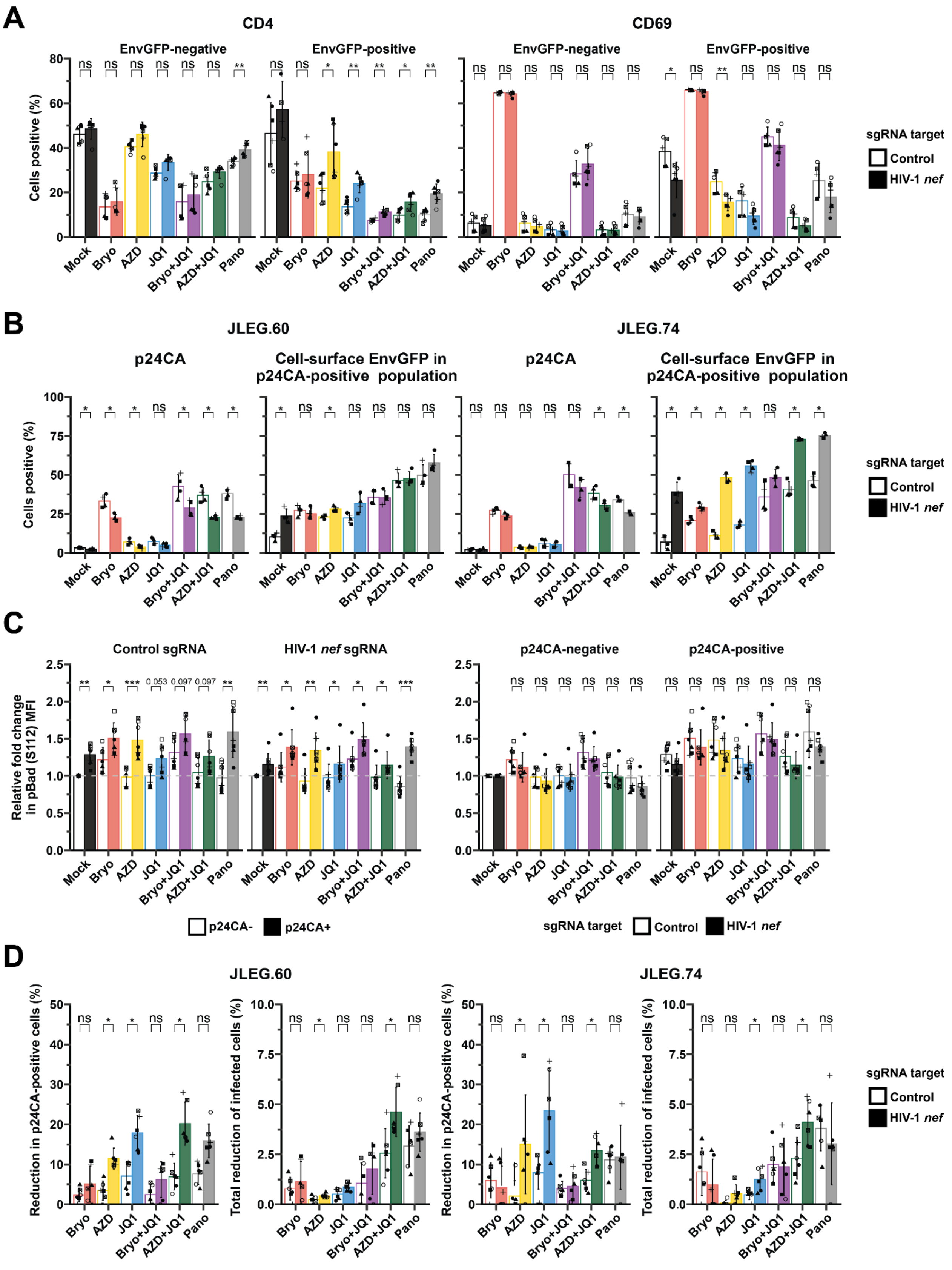

**Figure S8. HIV-1 *nef* knockout improves ADCC elimination efficiency in JLEG.60 and JLEG.74 clones.**

A) Cell-surface expression of CD4, CD69, HLA-E and PVR in *nef* knockout and control JLEG.1 T-cells for EnvGFP-negative and EnvGFP-positive cells in the same sample. measured by flow cytometry; n=6.

B) Expression of p24 and cell-surface EnvGFP in the p24CA-positive population of *nef* knockout and control JLEG.60 and JLEG.74 T-cells, either mock-treated or treated with the indicated LRAs for 24h, measured by flow cytometry; n=6.

C) Expression of pBad S112 in in *nef* knock-out and control JLEG.1 T-cells for p24CA-negative and p24CA-positive cells in the same sample, either mock-treated or treated with the indicated LRAs for 24h, measured by flow cytometry; n=6. The same data is presented in the top and bottom row, but rearranged to allow for clear indication of which statistical comparisons were made.

D) ADCC killing efficiency (left) and elimination efficiency (right) for *nef* knockout and control JLEG.60 and JLEG.74 T-cells, treated as in A) and co-cultured with PBMCs from HIV-negative donors in the presence of sera from sera from PLWH for 4h to induce cytotoxic killing. The ADCC killing efficiency was normalised to the level of p24CA-positive cells induced by each treatment to yield the elimination efficiency; n=6 donors.

Wilcoxon rank-signed test. *** $p < 0.001$; ** $p < 0.01$; * $p< 0.05$. Indicated is the arithmetic mean ± SD.

### REFERENCES

1. Brink, D.T., Martin-Hughes, R., Bowring, A.L., Wulan, N., Burke, K., Tidhar, T., Dalal, S., and Scott, N. (2025). Impact of an international HIV funding crisis on HIV infections and mortality in low-income and middle-income countries: a modelling study. Lancet HIV *0*. https://doi.org/10.1016/s2352-3018(25)00074-8.

2. Gruell, H., Gunst, J.D., Cohen, Y.Z., Pahus, M.H., Malin, J.J., Platten, M., Millard, K.G., Tolstrup, M., Jones, R.B., Conce Alberto, W.D., et al. (2022). Effect of 3BNC117 and romidepsin on the HIV-1 reservoir in people taking suppressive antiretroviral therapy (ROADMAP): a randomised, open-label, phase 2A trial. Lancet Microbe *3*, e203–e214.

3. Gunst, J.D., Pahus, M.H., Rosás-Umbert, M., Lu, I.-N., Benfield, T., Nielsen, H., Johansen, I.S., Mohey, R., Østergaard, L., Klastrup, V., et al. (2022). Early intervention with 3BNC117 and romidepsin at antiretroviral treatment initiation in people with HIV-1: a phase 1b/2a, randomized trial. Nat. Med. *28*, 2424–2435.

4. Nixon, C.C., Mavigner, M., Sampey, G.C., Brooks, A.D., Spagnuolo, R.A., Irlbeck, D.M., Mattingly, C., Ho, P.T., Schoof, N., Cammon, C.G., et al. (2020). Systemic HIV and SIV latency reversal via non-canonical NF-κB signalling in vivo. Nature *578*, 160–165.

5. Li, G., Zhang, Z., Reszka-Blanco, N., Li, F., Chi, L., Ma, J., Jeffrey, J., Cheng, L., and Su, L. (2019). Specific activation in vivo of HIV-1 by a bromodomain inhibitor from monocytic cells in humanized mice under antiretroviral therapy. J. Virol. *93*, e00233–19.

6. Dashti, A., Sukkestad, S., Horner, A.M., Neja, M., Siddiqi, Z., Waller, C., Goldy, J., Monroe, D., Lin, A., Schoof, N., et al. (2023). AZD5582 plus SIV-specific antibodies reduce lymph node viral reservoirs in antiretroviral therapy-suppressed macaques. Nat. Med. *29*, 2535–2546.

7. Debrabander, Q., Hensley, K.S., Psomas, C.K., Bramer, W., Mahmoudi, T., van Welzen, B.J., Verbon, A., and Rokx, C. (2023). The efficacy and tolerability of latency-reversing agents in reactivating the HIV-1 reservoir in clinical studies: a systematic review. J. Virus Erad. *9*, 100342.

8. Darcis, G., Kula, A., Bouchat, S., Fujinaga, K., Corazza, F., Ait-Ammar, A., Delacourt, N., Melard, A., Kabeya, K., Vanhulle, C., et al. (2015). An In-Depth Comparison of Latency-Reversing Agent Combinations in Various In Vitro and Ex Vivo HIV-1 Latency Models Identified Bryostatin-1+JQ1 and Ingenol-B+JQ1 to Potently Reactivate Viral Gene Expression. PLoS Pathog. *11*. https://doi.org/10.1371/JOURNAL.PPAT.1005063.

9. Falcinelli, S.D., Peterson, J.J., Turner, A.-M.W., Irlbeck, D., Read, J., Raines, S.L., James, K.S., Sutton, C., Sanchez, A., Emery, A., et al. (2022). Combined noncanonical NF-κB agonism and targeted BET bromodomain inhibition reverse HIV latency ex vivo. J. Clin. Invest. *132*. https://doi.org/10.1172/JCI157281.

10. Rasmussen, T.A., Tolstrup, M., Brinkmann, C.R., Olesen, R., Erikstrup, C., Solomon, A., Winckelmann, A., Palmer, S., Dinarello, C., Buzon, M., et al. (2014). Panobinostat, a histone deacetylase inhibitor, for latent-virus reactivation in HIV-infected patients on suppressive antiretroviral therapy: a phase 1/2, single group, clinical trial. Lancet HIV *1*, e13–e21.

11. Laird, G.M., Bullen, C.K., Rosenbloom, D.I.S., Martin, A.R., Hill, A.L., Durand, C.M., Siliciano, J.D., and Siliciano, R.F. (2015). Ex vivo analysis identifies effective HIV-1 latency-reversing drug combinations. J. Clin. Invest. *125*, 1901–1912.

12. Nakane, S., Iwamoto, A., and Matsuda, Z. (2015). The V4 and V5 Variable Loops of HIV-1 Envelope Glycoprotein Are Tolerant to Insertion of Green Fluorescent Protein and Are Useful Targets for Labeling. J. Biol. Chem. *290*, 15279.

13. Milacic, M., Beavers, D., Conley, P., Gong, C., Gillespie, M., Griss, J., Haw, R., Jassal, B., Matthews, L., May, B., et al. (2024). The reactome pathway knowledgebase 2024. Nucleic Acids Res. *52*, D672–D678.

14. van 't Wout, A.B., Swain, J.V., Schindler, M., Rao, U., Pathmajeyan, M.S., Mullins, J.I., and Kirchhoff, F. (2005). Nef induces multiple genes involved in cholesterol synthesis and uptake in human immunodeficiency virus type 1-infected T cells. J. Virol. *79*, 10053–10058.

15. Zheng, Y.-H., Plemenitas, A., Fielding, C.J., and Peterlin, B.M. (2003). Nef increases the synthesis of and transports cholesterol to lipid rafts and HIV-1 progeny virions. Proc. Natl. Acad. Sci. U. S. A. *100*, 8460–8465.

16. Cheng, Y., Jung, J., Guo, L., Shuboni-Mulligan, D.D., Chen, J.-F., Hu, W., and Guo, M.-L. (2025). HIV-TAT dysregulates microglial lipid metabolism through SREBP2/miR-124 axis: Implication of lipid droplet accumulation microglia in NeuroHIV. Brain Behav. Immun. *123*, 108–122.

17. Santis, A.G., Campanero, M.R., Alonso, J.L., Tugores, A., Alonso, M.A., Yagüe, E., Pivel, J.P., and Sánchez-Madrid, F. (1992). Tumor necrosis factor-alpha production induced in T lymphocytes through the AIM/CD69 activation pathway. Eur. J. Immunol. *22*, 1253–1259.

18. Thèze, J., Alzari, P.M., and Bertoglio, J. (1996). Interleukin 2 and its receptors: recent advances and new immunological functions. Immunol. Today *17*, 481–486.

19. Krapp, C., Hotter, D., Gawanbacht, A., McLaren, P.J., Kluge, S.F., Stürzel, C.M., Mack, K., Reith, E., Engelhart, S., Ciuffi, A., et al. (2016). Guanylate Binding Protein (GBP) 5 Is an Interferon-Inducible Inhibitor of HIV-1 Infectivity. Cell Host Microbe *19*, 504–514.

20. Braun, E., Hotter, D., Koepke, L., Zech, F., Groß, R., Sparrer, K.M.J., Müller, J.A., Pfaller, C.K., Heusinger, E., Wombacher, R., et al. (2019). Guanylate-binding proteins 2 and 5 exert broad antiviral activity by inhibiting furin-mediated processing of viral envelope proteins. Cell Rep. *27*, 2092–2104.e10.

21. Juneja, V.R., McGuire, K.A., Manguso, R.T., LaFleur, M.W., Collins, N., Haining, W.N., Freeman, G.J., and Sharpe, A.H. (2017). PD-L1 on tumor cells is sufficient for immune evasion in immunogenic tumors and inhibits CD8 T cell cytotoxicity. J. Exp. Med. *214*, 895–904.

22. Mehla, R., Bivalkar-Mehla, S., Zhang, R., Handy, I., Albrecht, H., Giri, S., Nagarkatti, P., Nagarkatti, M., and Chauhan, A. (2010). Bryostatin modulates latent HIV-1 infection via PKC and AMPK signaling but inhibits acute infection in a receptor independent manner. PLoS One *5*, e11160.

23. Hany, L., Turmel, M.-O., Barat, C., Ouellet, M., and Tremblay, M.J. (2022). Bryostatin-1

decreases HIV-1 infection and viral production in human primary macrophages. J. Virol. *96*, e0195321.

24. Grishkan, I.V., Ntranos, A., Calabresi, P.A., and Gocke, A.R. (2013). Helper T cells down-regulate CD4 expression upon chronic stimulation giving rise to double-negative T cells. Cell. Immunol. *284*, 68–74.

25. Lundquist, C.A., Tobiume, M., Zhou, J., Unutmaz, D., and Aiken, C. (2002). Nef-mediated downregulation of CD4 enhances human immunodeficiency virus type 1 replication in primary T lymphocytes. J. Virol. *76*, 4625–4633.

26. Hanna, Z., Priceputu, E., Hu, C., Vincent, P., and Jolicoeur, P. (2006). HIV-1 Nef mutations abrogating downregulation of CD4 affect other Nef functions and show reduced pathogenicity in transgenic mice. Virology *346*, 40–52.

27. daSilva, L.L.P., Sougrat, R., Burgos, P.V., Janvier, K., Mattera, R., and Bonifacino, J.S. (2009). Human immunodeficiency virus type 1 Nef protein targets CD4 to the multivesicular body pathway. J. Virol. *83*, 6578–6590.

28. Andrew, A., and Strebel, K. (2010). HIV-1 Vpu targets cell surface markers CD4 and BST-2 through distinct mechanisms. Mol. Aspects Med. *31*, 407–417.

29. Umviligihozo, G., Cobarrubias, K.D., Chandrarathna, S., Jin, S.W., Reddy, N., Byakwaga, H., Muzoora, C., Bwana, M.B., Lee, G.Q., Hunt, P.W., et al. (2020). Differential Vpu-mediated CD4 and tetherin downregulation functions among major HIV-1 group M subtypes. J. Virol. *94*. https://doi.org/10.1128/JVI.00293-20.

30. Michel, N., Allespach, I., Venzke, S., Fackler, O.T., and Keppler, O.T. (2005). The Nef protein of human immunodeficiency virus establishes superinfection immunity by a dual strategy to downregulate cell-surface CCR5 and CD4. Curr. Biol. *15*, 714–723.

31. Toyoda, M., Ogata, Y., Mahiti, M., Maeda, Y., Kuang, X.T., Miura, T., Jessen, H., Walker, B.D., Brockman, M.A., Brumme, Z.L., et al. (2015). Differential ability of primary HIV-1 Nef isolates to downregulate HIV-1 entry receptors. J. Virol. *89*, 9639–9652.

32. Venzke, S., Michel, N., Allespach, I., Fackler, O.T., and Keppler, O.T. (2006). Expression of Nef downregulates CXCR4, the major coreceptor of human immunodeficiency virus, from the surfaces of target cells and thereby enhances resistance to superinfection. J. Virol. *80*, 11141–11152.

33. Cai, C., Rodepeter, F.R., Rossmann, A., Teymoortash, A., Lee, J.-S., Quint, K., Dl Fazio, P., Ocker, M., Werner, J.A., and Mandic, R. (2012). SIVmac$_{239}$-Nef down-regulates cell surface expression of CXCR4 in tumor cells and inhibits proliferation, migration and angiogenesis. Anticancer Res. *32*, 2759–2768.

34. Veler, H., Lun, C.M., Waheed, A.A., and Freed, E.O. (2024). Guanylate-binding protein 5 antagonizes viral glycoproteins independently of furin processing. MBio *15*, e0208624.

35. Zahavi, D., AlDeghaither, D., O'Connell, A., and Weiner, L.M. (2018). Enhancing antibody-dependent cell-mediated cytotoxicity: a strategy for improving antibody-based immunotherapy. Antib. Ther. *1*, 7–12.

36. Matsuda, K., Islam, S., Takada, T., Tsuchiya, K., Yang Tan, B.J., Hattori, S.I., Katsuya, H., Kitagawa, K., Kim, K.S., Matsuo, M., et al. (2021). A widely distributed HIV-1 provirus elimination assay to evaluate latency-reversing agents in vitro. Cell Reports Methods *1*, 100122.

37. French, A.J., Natesampillai, S., Krogman, A., Correia, C., Peterson, K.L., Alto, A., Chandrasekar, A.P., Misra, A., Li, Y., Kaufmann, S.H., et al. (2020). Reactivating latent HIV with PKC agonists induces resistance to apoptosis and is associated with phosphorylation and activation of BCL2. PLoS Pathog. *16*. https://doi.org/10.1371/JOURNAL.PPAT.1008906.

38. Zha, J., Harada, H., Yang, E., Jockel, J., and Korsmeyer, S.J. (1996). Serine phosphorylation of death agonist BAD in response to survival factor results in binding to 14-3-3 not BCL-X(L). Cell *87*, 619–628.

39. Lee, N., Llano, M., Carretero, M., Ishitani, A., Navarro, F., López-Botet, M., and Geraghty, D.E. (1998). HLA-E is a major ligand for the natural killer inhibitory receptor CD94/NKG2A. Proc. Natl. Acad. Sci. U. S. A. *95*, 5199–5204.

40. Wolf, D., Witte, V., Laffert, B., Blume, K., Stromer, E., Trapp, S., d’Aloja, P., Schürmann, A., and Baur, A.S. (2001). HIV-1 Nef associated PAK and PI3-kinases stimulate Akt-independent Bad-phosphorylation to induce anti-apoptotic signals. Nat. Med. *7*, 1217–1224.

41. Perica, K., Kotchetkov, I.S., Mansilla-Soto, J., Ehrich, F., Herrera, K., Shi, Y., Dobrin, A., Gönen, M., and Sadelain, M. (2025). HIV immune evasin Nef enhances allogeneic CAR T cell potency. Nature *640*, 793–801.

42. Kumar, B., Tripathi, C., Kanchan, R.K., Tripathi, J.K., Ghosh, J.K., Ramachandran, R., Bhadauria, S., and Tripathi, R.K. (2013). Dynamics of physical interaction between HIV-1 Nef and ASK1: identifying the interacting motif(s). PLoS One *8*, e67586.

43. James, C.O., Huang, M.-B., Khan, M., Garcia-Barrio, M., Powell, M.D., and Bond, V.C. (2004). Extracellular Nef protein targets CD4+ T cells for apoptosis by interacting with CXCR4 surface receptors. J. Virol. *78*, 3099–3109.

44. Lenassi, M., Cagney, G., Liao, M., Vaupotic, T., Bartholomeeusen, K., Cheng, Y., Krogan, N.J., Plemenitas, A., and Peterlin, B.M. (2010). HIV Nef is secreted in exosomes and triggers apoptosis in bystander CD4+ T cells. Traffic *11*, 110–122.

45. Blagoveshchenskaya, A.D., Thomas, L., Feliciangeli, S.F., Hung, C.H., and Thomas, G. (2002). HIV-1 Nef downregulates MHC-I by a PACS-1- and PI3K-regulated ARF6 endocytic pathway. Cell *111*, 853–866.

46. Garcia, J.V., and Miller, A.D. (1991). Serine phosphorylation-independent downregulation of cell-surface CD4 by nef. Nature *350*, 508–511.

47. Matusali, G., Potestà, M., Santoni, A., Cerboni, C., and Doria, M. (2012). The human immunodeficiency virus type 1 Nef and Vpu proteins downregulate the natural killer cell-activating ligand PVR. J. Virol. *86*, 4496–4504.

48. van Stigt Thans, T., Akko, J.I., Niehrs, A., Garcia-Beltran, W.F., Richert, L., Stürzel, C.M., Ford, C.T., Li, H., Ochsenbauer, C., Kappes, J.C., et al. (2019). Primary HIV-1 strains use

Nef to downmodulate HLA-E surface expression. J. Virol. *93*, e00719–19.

49. Dieudonné, M., Maiuri, P., Biancotto, C., Knezevich, A., Kula, A., Lusic, M., and Marcello, A. (2009). Transcriptional competence of the integrated HIV-1 provirus at the nuclear periphery. EMBO J. *28*, 2231–2243.

50. Macedo, A.B., Novis, C.L., De Assis, C.M., Sorensen, E.S., Moszczynski, P., Huang, S.H., Ren, Y., Spivak, A.M., Jones, R.B., Planelles, V., et al. (2018). Dual TLR2 and TLR7 agonists as HIV latency-reversing agents. JCI insight *3*. https://doi.org/10.1172/jci.insight.122673.

51. Moran, J.A., Ranjan, A., Hourani, R., Kim, J.T., Wender, P.A., Zack, J.A., and Marsden, M.D. (2023). Secreted factors induced by PKC modulators do not indirectly cause HIV latency reversal. Virology *581*, 8–14.

52. Kim, E.H., Manganaro, L., Schotsaert, M., Brown, B.D., Mulder, L.C.F., and Simon, V. (2022). Development of an HIV reporter virus that identifies latently infected CD4+ T cells. Cell Rep. Methods *2*, 100238.

53. Cai, J., Gao, H., Zhao, J., Hu, S., Liang, X., Yang, Y., Dai, Z., Hong, Z., and Deng, K. (2021). Infection with a newly designed dual fluorescent reporter HIV-1 effectively identifies latently infected CD4+ T cells. Elife *10*. https://doi.org/10.7554/eLife.63810.

54. Dahabieh, M.S., Ooms, M., Simon, V., and Sadowski, I. (2013). A doubly fluorescent HIV-1 reporter shows that the majority of integrated HIV-1 is latent shortly after infection. J. Virol. *87*, 4716–4727.

55. Kok, Y.L., Schmutz, S., Inderbitzin, A., Neumann, K., Kelley, A., Jörimann, L., Shilaih, M., Vongrad, V., Kouyos, R.D., Günthard, H.F., et al. (2018). Spontaneous reactivation of latent HIV-1 promoters is linked to the cell cycle as revealed by a genetic-insulators-containing dual-fluorescence HIV-1-based vector. Sci. Rep. *8*, 10204.

56. Lu, Y., Singh, H., Singh, A., and Dar, R.D. (2021). A transient heritable memory regulates HIV reactivation from latency. iScience *24*, 102291.

57. Kim, Y., Anderson, J.L., and Lewin, S.R. (2018). Getting the “Kill” into “Shock and Kill”: Strategies to Eliminate Latent HIV. Preprint at Cell Press, https://doi.org/10.1016/j.chom.2017.12.004 https://doi.org/10.1016/j.chom.2017.12.004.

58. Crooks, A.M., Bateson, R., Cope, A.B., Dahl, N.P., Griggs, M.K., Kuruc, J.D., Gay, C.L., Eron, J.J., Margolis, D.M., Bosch, R.J., et al. (2015). Precise quantitation of the latent HIV-1 reservoir: Implications for eradication strategies. J. Infect. Dis. *212*, 1361–1365.

59. Cevaal, P.M., Kan, S., Fisher, B.M., Moso, M.A., Tan, A., Liu, H., Ali, A., Tanaka, K., Shepherd, R.A., Kim, Y., et al. (2025). Efficient mRNA delivery to resting T cells to reverse HIV latency. Nat. Commun. *16*, 4979.

60. Timmons, A., Fray, E., Kumar, M., Wu, F., Dai, W., Bullen, C.K., Kim, P., Hetzel, C., Yang, C., Beg, S., et al. (2020). HSF1 inhibition attenuates HIV-1 latency reversal mediated by several candidate LRAs In Vitro and Ex Vivo. Proc. Natl. Acad. Sci. U. S. A. *117*, 15763–15771.

61. Rodari, A., Darcis, G., and Van Lint, C.M. (2021). The current status of latency reversing agents for HIV-1 remission. Annu. Rev. Virol. *8*, 491–514.

62. Heylmann, D., Badura, J., Becker, H., Fahrer, J., and Kaina, B. (2018). Sensitivity of CD3/CD28-stimulated versus non-stimulated lymphocytes to ionizing radiation and genotoxic anticancer drugs: key role of ATM in the differential radiation response. Cell Death Dis. *9*, 1053.

63. Su, C.-C., Lin, Y.-P., Cheng, Y.-J., Huang, J.-Y., Chuang, W.-J., Shan, Y.-S., and Yang, B.-C. (2007). Phosphatidylinositol 3-kinase/Akt activation by integrin-tumor matrix interaction suppresses Fas-mediated apoptosis in T cells. J. Immunol. *179*, 4589–4597.

64. Karas, M., Zaks, T.Z., Yakar, S., Dudley, M.E., and LeRoith, D. (2001). TCR stimulation protects CD8+ T cells from CD95 mediated apoptosis. Hum. Immunol. *62*, 32–38.

65. Varadhachary, A.S., Perdow, S.N., Hu, C., Ramanarayanan, M., and Salgame, P. (1997). Differential ability of T cell subsets to undergo activation-induced cell death. Proc. Natl. Acad. Sci. U. S. A. *94*, 5778–5783.

66. Meier, A., Bagchi, A., Sidhu, H.K., Alter, G., Suscovich, T.J., Kavanagh, D.G., Streeck, H., Brockman, M.A., LeGall, S., Hellman, J., et al. (2008). Upregulation of PD-L1 on monocytes and dendritic cells by HIV-1 derived TLR ligands. AIDS *22*, 655–658.

67. Boasso, A., Hardy, A.W., Landay, A.L., Martinson, J.L., Anderson, S.A., Dolan, M.J., Clerici, M., and Shearer, G.M. (2008). PDL-1 upregulation on monocytes and T cells by HIV via type I interferon: restricted expression of type I interferon receptor by CCR5-expressing leukocytes. Clin. Immunol. *129*, 132–144.

68. Hess, A.D., Silanskis, M.K., Esa, A.H., Pettit, G.R., and May, W.S. (1988). Activation of human T lymphocytes by bryostatin. J. Immunol. *141*, 3263–3269.

69. Esa, A.H., Boto, W.O., Adler, W.H., May, W.S., and Hess, A.D. (1990). Activation of T-cells by bryostatins: induction of the IL-2 receptor gene transcription and down-modulation of surface receptors. Int. J. Immunopharmacol. *12*, 481–490.

70. Richard, J., Prévost, J., Alsahafi, N., Ding, S., and Finzi, A. (2018). Impact of HIV-1 envelope conformation on ADCC responses. Trends Microbiol. *26*, 253–265.

71. Mielke, D., Bandawe, G., Zheng, J., Jones, J., Abrahams, M.-R., Bekker, V., Ochsenbauer, C., Garrett, N., Abdool Karim, S., Moore, P.L., et al. (2021). ADCC-mediating non-neutralizing antibodies can exert immune pressure in early HIV-1 infection. PLoS Pathog. *17*, e1010046.

72. Arandjelovic, P., Kim, Y., Cooney, J.P., Preston, S.P., Doerflinger, M., McMahon, J.H., Garner, S.E., Zerbato, J.M., Roche, M., Tumpach, C., et al. (2023). Venetoclax, alone and in combination with the BH3 mimetic S63845, depletes HIV-1 latently infected cells and delays rebound in humanized mice. Cell Rep. Med. *4*, 101178.

73. Alto, A., Natesampillai, S., Chandrasekar, A.P., Krogman, A., Misra, A., Shweta, F.N.U., VanLith, C., Yao, J.D., Cummins, N.W., and Badley, A.D. (2021). The Combination of Venetoclax and Ixazomib Selectively and Efficiently Kills HIV-Infected Cell Lines but Has Unacceptable Toxicity in Primary Cell Models. J. Virol. *95*.

https://doi.org/10.1128/JVI.00138-21.

74. Klinnert, S., Schenkel, C.D., Freitag, P.C., Günthard, H.F., Plückthun, A., and Metzner, K.J. (2024). Targeted shock-and-kill HIV-1 gene therapy approach combining CRISPR activation, suicide gene tBid and retargeted adenovirus delivery. Gene Ther. *31*, 74–84.

75. Moon, J.-H., Shin, J.-S., Hong, S.-W., Jung, S.-A., Hwang, I.-Y., Kim, J.H., Choi, E.K., Ha, S.-H., Kim, J.-S., Kim, K.-M., et al. (2015). A novel small-molecule IAP antagonist, AZD5582, draws Mcl-1 down-regulation for induction of apoptosis through targeting of cIAP1 and XIAP in human pancreatic cancer. Oncotarget *6*, 26895–26908.

76. Harada, H., Becknell, B., Wilm, M., Mann, M., Huang, L.J., Taylor, S.S., Scott, J.D., and Korsmeyer, S.J. (1999). Phosphorylation and inactivation of BAD by mitochondria-anchored protein kinase A. Mol. Cell *3*, 413–422.

77. Mok, C.L., Gil-Gómez, G., Williams, O., Coles, M., Taga, S., Tolaini, M., Norton, T., Kioussis, D., and Brady, H.J. (1999). Bad can act as a key regulator of T cell apoptosis and T cell development. J. Exp. Med. *189*, 575–586.

78. She, Q.-B., Solit, D.B., Ye, Q., O'Reilly, K.E., Lobo, J., and Rosen, N. (2005). The BAD protein integrates survival signaling by EGFR/MAPK and PI3K/Akt kinase pathways in PTEN-deficient tumor cells. Cancer Cell *8*, 287–297.

79. Kane, L.P., and Weiss, A. (2003). The PI-3 kinase/Akt pathway and T cell activation: pleiotropic pathways downstream of PIP3: Kane & Weiss · PI-3 kinase, Akt, and T cell activation. Immunol. Rev. *192*, 7–20.

80. Herrero-Sánchez, M.C., Rodríguez-Serrano, C., Almeida, J., San Segundo, L., Inogés, S., Santos-Briz, Á., García-Briñón, J., Corchete, L.A., San Miguel, J.F., Del Cañizo, C., et al. (2016). Targeting of PI3K/AKT/mTOR pathway to inhibit T cell activation and prevent graft-versus-host disease development. J. Hematol. Oncol. *9*, 113.

81. Xue, L., Chiang, L., Kang, C., and Winoto, A. (2008). The role of the PI3K-AKT kinase pathway in T-cell development beyond the beta checkpoint. Eur. J. Immunol. *38*, 3200–3207.

82. D'Souza, W.N., Chang, C.-F., Fischer, A.M., Li, M., and Hedrick, S.M. (2008). The Erk2 MAPK regulates CD8 T cell proliferation and survival. J. Immunol. *181*, 7617–7629.

83. Marquis, M., Boulet, S., Mathien, S., Rousseau, J., Thébault, P., Daudelin, J.-F., Rooney, J., Turgeon, B., Beauchamp, C., Meloche, S., et al. (2014). The non-classical MAP kinase ERK3 controls T cell activation. PLoS One *9*, e86681.

84. Prévost, J., Richard, J., Gasser, R., Medjahed, H., Kirchhoff, F., Hahn, B.H., Kappes, J.C., Ochsenbauer, C., Duerr, R., and Finzi, A. (2022). Detection of the HIV-1 accessory proteins Nef and Vpu by flow cytometry represents a new tool to study their functional interplay within a single infected CD4+ T cell. J. Virol. *96*, e0192921.

85. Meylan, P.R., Baumgartner, M., Ciuffi, A., Munoz, M., and Sahli, R. (1998). The nef gene controls syncytium formation in primary human lymphocytes and macrophages infected by HIV type 1. AIDS Res. Hum. Retroviruses *14*, 1531–1542.

86. Schwartz, O., Rivière, Y., Heard, J.M., and Danos, O. (1993). Reduced cell surface expression of processed human immunodeficiency virus type 1 envelope glycoprotein in the presence of Nef. J. Virol. *67*, 3274–3280.

87. Veillette, M., Coutu, M., Richard, J., Batraville, L.-A., Dagher, O., Bernard, N., Tremblay, C., Kaufmann, D.E., Roger, M., and Finzi, A. (2015). The HIV-1 gp120 CD4-bound conformation is preferentially targeted by antibody-dependent cellular cytotoxicity-mediating antibodies in sera from HIV-1-infected individuals. J. Virol. *89*, 545–551.

88. Prévost, J., Richard, J., Medjahed, H., Alexander, A., Jones, J., Kappes, J.C., Ochsenbauer, C., and Finzi, A. (2018). Incomplete downregulation of CD4 expression affects HIV-1 Env conformation and antibody-dependent cellular cytotoxicity responses. J. Virol. *92*. https://doi.org/10.1128/JVI.00484-18.

89. Dufloo, J., Guivel-Benhassine, F., Buchrieser, J., Lorin, V., Grzelak, L., Dupouy, E., Mestrallet, G., Bourdic, K., Lambotte, O., Mouquet, H., et al. (2020). Anti-HIV-1 antibodies trigger non-lytic complement deposition on infected cells. EMBO Rep. *21*, e49351.

90. Gutiérrez, C., Serrano-Villar, S., Madrid-Elena, N., Pérez-Elías, M.J., Martín, M.E., Barbas, C., Ruipérez, J., Muñoz, E., Muñoz-Fernández, M.A., Castor, T., et al. (2016). Bryostatin-1 for latent virus reactivation in HIV-infected patients on antiretroviral therapy. AIDS *30*, 1385–1392.

91. Prigann, J., Postmus, D., Pietrobon, A.J., Wyler, E., Jansen, J., Möller, L., Fadejeva, J., Steijaert, T.H., Fischer, C., Koppe, U., et al. (2020). Type I interferons sensitise HIV-1-reactivating T-cells for NK cell-mediated elimination despite HDACi-imposed dysregulation of innate immunity. Microbiology.

92. Brinkmann, C.R., Højen, J.F., Rasmussen, T.A., Kjær, A.S., Olesen, R., Denton, P.W., Østergaard, L., Ouyang, Z., Lichterfeld, M., Yu, X., et al. (2018). Treatment of HIV-Infected Individuals with the Histone Deacetylase Inhibitor Panobinostat Results in Increased Numbers of Regulatory T Cells and Limits Ex Vivo Lipopolysaccharide-Induced Inflammatory Responses. mSphere *3*. https://doi.org/10.1128/msphere.00616-17.

93. Dai, W., Wu, F., McMyn, N., Song, B., Walker-Sperling, V.E., Varriale, J., Zhang, H., Barouch, D.H., Siliciano, J.D., Li, W., et al. (2022). Genome-wide CRISPR screens identify combinations of candidate latency reversing agents for targeting the latent HIV-1 reservoir. Sci. Transl. Med. *14*, eabh3351.

94. Slavic Obradovic, K., Ebner, F., Artemov, A.V., Miotto, M., Traexler, P.-E., Jacob, R., Pham Thi Thanh, H., Ruzicka, R., Wernitznig, A., Baumann, I., et al. (2025). Combining BET inhibition with SMAC mimetics restricts tumor growth and triggers immune surveillance in preclinical cancer models. Cell Rep. Med. *6*, 102313.

95. Staudt, R.P., Alvarado, J.J., Emert-Sedlak, L.A., Shi, H., Shu, S.T., Wales, T.E., Engen, J.R., and Smithgall, T.E. (2020). Structure, function, and inhibitor targeting of HIV-1 Nef-effector kinase complexes. J. Biol. Chem. *295*, 15158.

96. Emert-Sedlak, L.A., Loughran, H.M., Shi, H., Kulp, J.L., 3rd, Shu, S.T., Zhao, J., Day, B.W., Wrobel, J.E., Reitz, A.B., and Smithgall, T.E. (2016). Synthesis and evaluation of orally active small molecule HIV-1 Nef antagonists. Bioorg. Med. Chem. Lett. *26*, 1480–1484.

97. Emert-Sedlak, L.A., Tice, C.M., Shi, H., Alvarado, J.J., Shu, S.T., Reitz, A.B., and Smithgall, T.E. (2024). PROTAC-mediated degradation of HIV-1 Nef efficiently restores cell-surface CD4 and MHC-I expression and blocks HIV-1 replication. Cell Chem. Biol. *31*, 658–668.e14.

98. Mujib, S., Saiyed, A., Fadel, S., Bozorgzad, A., Aidarus, N., Yue, F.Y., Benko, E., Kovacs, C., Emert-Sedlak, L.A., Smithgall, T.E., et al. (2017). Pharmacologic HIV-1 Nef blockade promotes CD8 T cell-mediated elimination of latently HIV-1-infected cells in vitro. JCI Insight *2*. https://doi.org/10.1172/jci.insight.93684.

99. Doench, J.G., Fusi, N., Sullender, M., Hegde, M., Vaimberg, E.W., Donovan, K.F., Smith, I., Tothova, Z., Wilen, C., Orchard, R., et al. (2016). Optimized sgRNA design to maximize activity and minimize off-target effects of CRISPR-Cas9. Nat. Biotechnol. *34*, 184–191.

100. Labun, K., Montague, T.G., Gagnon, J.A., Thyme, S.B., and Valen, E. (2016). CHOPCHOP v2: a web tool for the next generation of CRISPR genome engineering. Nucleic Acids Res. *44*, W272–W276.

101. Labun, K., Montague, T.G., Krause, M., Torres Cleuren, Y.N., Tjeldnes, H., and Valen, E. (2019). CHOPCHOP v3: expanding the CRISPR web toolbox beyond genome editing. Nucleic Acids Res. *47*, W171–W174.

102. Stewart, S.A., Dykxhoorn, D.M., Palliser, D., Mizuno, H., Yu, E.Y., An, D.S., Sabatini, D.M., Chen, I.S.Y., Hahn, W.C., Sharp, P.A., et al. (2003). Lentivirus-delivered stable gene silencing by RNAi in primary cells. RNA *9*, 493–501.

103. Liszewski, M.K., Yu, J.J., and O'Doherty, U. (2009). Detecting HIV-1 integration by repetitive-sampling Alu-gag PCR. Methods *47*, 254–260.

104. Cameron, D.L., Jacobs, N., Roepman, P., Priestley, P., Cuppen, E., and Papenfuss, A.T. (2021). VIRUSBreakend: Viral integration recognition using single breakends. Bioinformatics *37*, 3115–3119.

105. Cameron, D.L., Baber, J., Shale, C., Valle-Inclan, J.E., Besselink, N., van Hoeck, A., Janssen, R., Cuppen, E., Priestley, P., and Papenfuss, A.T. (2021). GRIDSS2: comprehensive characterisation of somatic structural variation using single breakend variants and structural variant phasing. Genome Biol. *22*, 202.

106. Danecek, P., Bonfield, J.K., Liddle, J., Marshall, J., Ohan, V., Pollard, M.O., Whitwham, A., Keane, T., McCarthy, S.A., Davies, R.M., et al. (2021). Twelve years of SAMtools and BCFtools. Gigascience *10*, giab008.

107. Dobin, A., Davis, C.A., Schlesinger, F., Drenkow, J., Zaleski, C., Jha, S., Batut, P., Chaisson, M., and Gingeras, T.R. (2013). STAR: ultrafast universal RNA-seq aligner. Bioinformatics *29*, 15–21.

108. Liao, Y., Smyth, G.K., and Shi, W. (2019). The R package Rsubread is easier, faster, cheaper and better for alignment and quantification of RNA sequencing reads. Nucleic Acids Res. *47*, e47.

109. Love, M.I., Huber, W., and Anders, S. (2014). Moderated estimation of fold change and dispersion for RNA-seq data with DESeq2. Genome Biol. *15*, 550.

110. R Core Team The R Project for Statistical Computing. The R Project for Statistical Computing. https://www.r-project.org/.

111. Wickham, H. (2016). ggplot2: Elegant Graphics for Data Analysis. Preprint at Springer-Verlag New York.

112. Kolde, R. (2025). pheatmap: Pretty Heatmaps. Preprint.

113. Kassambara, A. (2023). rstatix: Pipe-Friendly Framework for Basic Statistical Tests. Preprint.

114. Van der Borght, K., Tourny, A., Bagdziunas, R., Thas, O., Nazarov, M., Turner, H., Verbist, B., and Ceulemans, H. (2017). BIGL: Biochemically Intuitive Generalized Loewe null model for prediction of the expected combined effect compatible with partial agonism and antagonism. Sci. Rep. 7, 17935.
